# Tissue-specific regulation of gene expression via unproductive splicing

**DOI:** 10.1101/2022.07.03.498634

**Authors:** Alexey Mironov, Marina Petrova, Sergei Margasyuk, Maria Vlasenok, Andrei A. Mironov, Dmitry Skvortsov, Dmitri D. Pervouchine

**Affiliations:** Skolkovo Institute of Science and Technology, Center of Molecular and Cellular Biology, Bolshoy blv. 30, Moscow 121205, Russia; Moscow State University, Faculty of Bioengineering and Bioinformatics, ul. Kolmogorova 1, Moscow 119991, Russia; Moscow State University, Faculty of Chemistry, ul. Kolmogorova 1, Moscow 119991, Russia

## Abstract

Eukaryotic gene expression is regulated post-transcriptionally by a mechanism called unproductive splicing, in which mRNA is triggered to degradation by the nonsense-mediated decay (NMD) pathway as a result of regulated alternative splicing (AS). Only a few dozen unproductive splicing events (USEs) are currently documented, and many more remain to be identified. Here, we analyzed RNA-seq experiments from the Genotype-Tissue Expression (GTEx) Consortium to identify USEs, in which an increase in the NMD isoform splicing rate is accompanied by tissue-specific downregulation of the host gene. Further, to characterize RBPs that regulate USEs, we superimposed these results with RNA-binding protein (RBP) footprinting data and experiments on the response of the transcriptome to the perturbation of expression of a large panel of RBPs. Concordant tissue-specific changes between the expression of RBP and USE splicing rate revealed a high-confidence regulatory network including 27 tissue-specific USEs with strong evidence of RBP binding. Among them, we found previously unknown PTBP1-controlled events in the DCLK2 and IQGAP1 genes, for which we confirmed the regulatory effect using siRNA-knockdown experiments in the A549 cell line. In sum, we present a transcriptomic pipeline that allows the identification of tissue-specific USEs, potentially many more than we have reported here using stringent filters.

## Introduction

Nonsense mutations and frame-shifting splicing errors give rise to transcripts with premature termination codons (PTC), which encode truncated, deleterious proteins. In eukaryotes, such transcripts are selectively degraded by a translation-dependent mRNA surveillance pathway called the Nonsense-Mediated mRNA Decay (NMD) [1]. However, NMD serves not only as mRNA quality control system but also as a regulatory homeostatic mechanism to maintain the abundance of a broad class of physiological transcripts [2, 3, 4, 5, 6]. In this mechanism, referred to as regulated unproductive splicing and translation (RUST) [7, 8] or simply unproductive splicing [9, 10], the cell employs alternative splicing (AS) to produce a PTC-containing transcript in order to post-transcriptionally downregulate the expression level of the gene [11, 9]. Unproductive splicing plays an essential role in development and disease including early embryogenesis [4], granulocyte differentiation [12], and stabilization or repression of oncogenic expression [13, 14].

The majority of unproductive splicing events (USEs) are described in genes encoding RNA-binding proteins (RBPs) [15]. Many of them use unproductive splicing to autoregulate their expression levels in a negative feedback loop, in which the gene product binds its own pre-mRNA and causes alternative splicing to induce a PTC. This autoregulation takes place in almost all serine-arginine-rich (SR) proteins [16], many hnRNP proteins [17, 18, 19, 20], spliceosome components [21, 22], and even in ribosomal proteins [23, 24]. Autoregulatory unproductive splicing is found in almost all eukaryotes studied to date and exhibits a high degree of evolutionary conservation [25, 26, 27].

Cross-regulatory unproductive splicing networks have a different hierarchical organization compared to transcriptional networks, with much fewer master regulators and many more regulatory connections among RBPs than between RBPs and other genes [8]. These connections have been characterized for many splicing factors, particularly for SR proteins, which coordinate their expression in a dense unproductive splicing network [16]. Cross-regulatory circuits among paralogs such as PTBP1/PTBP2 [28, 17], SRSF3/SRSF7 [29, 30], RBM10/RBM5 [21], RBFOX2/RBFOX3 [31], hnRNPD/ hnRNPDL [20], and hnRNPL/hnRNPLL [32] are particularly abundant, but unproductive splicing also extends beyond RBPs and shapes the transcriptional landscape in other gene classes [15]. Remarkably, the relationship between the expression of the NMD isoform and the mRNA or protein levels in many cases is complex due to indirect connectivities in the regulatory network [19, 33, 34, 35]. Tissue specificity of unproductive splicing has been studied only for a handful of cases [15], including the regulation of neural-specific expression of the postsynaptic proteins DLG4 and GABBR1 that is controlled by PTBP1 and PTBP2 [36, 37, 38], which are also discussed here.

IIn this work, we interrogated tissue-specific unproductive splicing using a large panel of human tissue transcriptomes from the Genotype-Tissue Expression (GTEx) project [39]. Toward this goal, we cataloged USEs by extracting AS events that generate NMD isoforms, identified USEs with a significant negative association between gene expression level and NMD isoform splicing rate, and characterized tissue-specificity of this association. Further, to identify RBPs that regulate USEs, we superimposed these results with RBP footprinting data from the POSTAR3 database [40] and publicly available data on the transcriptome response to RBP expression perturbations, including experiments by ENCODE Consortium [41]. Finally, we identified USEs that responded significantly to RBP inactivation and also had concordant changes of RBP expression and USE splicing rate in GTEx tissues. This analysis yielded a high-confidence regulatory network of 27 tissue-specific USEs with strong evidence of RBP binding, from which we selected two prominent PTBP1 targets, DCLK2 and IQGAP1, and confirmed their unproductive splicing regulation using siRNA-mediated knockdown of *PTBP1* in the A549 cell line.

## MATERIALS AND METHODS

### Unproductive splicing events

The GRCh37 (hg19) assembly of the human genome was downloaded from the UCSC genome browser website [42]. It was used along with comprehensive gene annotation (v35lift37) from the GENCODE Consortium [43]. The coordinates of exons, introns, and stop codons were extracted from transcripts labeled as protein-coding and NMD to identify AS events shown in Figure S1. Among them, USEs were characterized by PTCs that were located within or downstream of the AS event, with the exception of poison exons and intron detention, which can trigger NMD by inducing splice junctions in the 3’-UTR. As a result, an initial set of 5,309 USEs was obtained. It was merged with the curated list of validated USEs obtained by literature search (Table S1) resulting in a total of 5322 USEs (Table S2). As a control, we used a set of 38,396 AS events that generated protein-coding isoforms according to GENCODE annotation.

### Quantification of AS

STAR v2.4.2a alignments [44] of short reads in 8551 samples from the Genotype-Tissue Expression (GTEx) Consortium v7 [45] (excluding samples from transformed cells and testis) were obtained from dbGaP website in BAM format under the accession number phs000424/GRU. Split reads supporting splice junctions as well as non-split reads supporting intron retention were extracted from BAM files using the IPSA package with the default settings [46].

Iin the GTEx samples, the splicing rate was estimated from split-read counts computed by the IPSA pipeline using the *ψ* (percent-spliced-in, PSI) metric, defined as the number of reads supporting the NMD isoform as a fraction of the combined number of reads supporting the NMD isoform and the protein-coding isoform (Figure S1). For uniformity, the *ψ* metric was always defined with respect to the NMD isoform, i.e., *ψ* = 1 for a poison exon assumes that it is 100% included, but *ψ* = 1 for an essential exon assumes that it is 100% skipped. Only *ψ* values with the denominator of at least 15 were considered. Further, we discarded AS events if the lower and the upper quartiles of the *ψ* distribution in GTEx samples coincided, or if *ψ* values were missing in more than 80% of samples of a tissue.

The results of RBP perturbation followed by RNA-seq (Table S3) including knockdown (KD), knockout (KO), and overexpression (OE) experiments and the results of siRNA-mediated PTBP1 KD with fractionation into cytoplasmic and nuclear RNA [47] were downloaded from the ENCODE portal and the SRA archive in BAM and FASTQ formats, respectively [41]. Short reads were mapped to the GRCh37 assembly of the human genome using STAR-2.7.7a aligner in two-pass mode with the following options: --outSJfilterOverhangMin 20 8 20 20 -- alignSJDBoverhangMin 8 --outFilterMismatchNmax 999 –outFilterMismatchNoverReadLmax 0.1. In differential splicing analysis, BAM files were processed with rMATS v.4.1.1 [48] with the following parameters: --novelSS --mil 10 --mel 10000 --libType fr-unstranded --variable-read-length. Only JC output files were used, i.e. differential splicing analysis did not involve reads mapped within exons.

### Gene expression quantification and analysis

The global gene expression level (*e*_*g*_) was defined as the total number of reads mapped to a gene that harbors the USE (i.e., read counts obtained from the GTEx portal). The local gene expression level (*e*_*l*_) was defined as the denominator of the *ψ* ratio of an USE, which reflects the abundance of short reads in its vicinity and allows estimation of gene expression level independently of AS acting globally in the gene. Both *e*_*g*_ and *e*_*l*_ values were normalized according to the DESeq2 methodology with a pseudocount of eight [49]. Namely, each row of the expression matrix, whose columns correspond to samples and rows correspond to genes, was divided by the row median. The size factor *s f*_*k*_ of the sample *k* was estimated as the median of the *k*-th column. Each gene expression value was divided by *s f*_*k*_ and log_2_-transformed. The resulting expression values were centered by subtracting the median over all samples.

To evaluate differential gene expression in RBP perturbation experiments, read counts were extracted from the respective BAM files using RNASeQC utility [50] and processed with DESeq2 package [49] using apeglm shrinkage correction [51]. In each RBP perturbation experiment, we checked that RBP expression changed upon perturbation in the intended direction, i.e. decreased upon KD or KO and increased upon OE; otherwise, the experiment was excluded from the analysis.

### Unproductive splicing and gene expression

To detect the downregulation of gene expression level that was accompanied by the activation of the NMD isoform, we compared gene expression levels in the 25% of GTEx samples with the highest *ψ* value (the upper quartile) vs. that in the 25% of GTEx samples with the lowest *ψ* value (the lower quartile) for each USE. Namely, we estimated the medians of *e*_*l*_ and *e*_*g*_ in these sample groups and compared them using the Mann-Whitney sum of ranks test with continuity correction. The statistical significance of the differences in *e*_*l*_ and *e*_*g*_ between the two groups (denoted as ∈*e*_*l*_ and ∈*e*_*g*_) was interpreted by *z*-scores associated with the U-statistic of the test.

Next, we determined the features that distinguish USEs from protein-coding AS events. First, we selected USEs, in which both ∈*e*_*l*_ and ∈*e*_*g*_ were positive (positive set) and USEs, in which both ∈*e*_*l*_ and ∈*e*_*g*_ were negative (negative set). In most AS types, the negative set was substantially larger among USEs compared to protein-coding AS events (Figure S2A) indicating that the negative association between AS and the host gene expression is more prevalent in USEs. Next, we compared the distributions of five parameters in the negative set for USEs and protein-coding AS events using the positive set as a control (Figure S2B-I). We found that the most remarkable feature that distinguished USEs and protein-coding AS events was the *z*-score of ∈*e*_*g*_. We checked additionally that gene expression levels don’t show a significant dependence on *ψ* value for AS events in protein-coding genes (Figure S3). In the downstream analysis, we required that significant changes of *e*_*g*_ co-occur with significant changes of *e*_*l*_.

### Tissue specificity of USEs

To assess tissue specificity of an USE, we tested the significance of deviations of *ψ, e*_*l*_, and *e*_*g*_ from their respective medians across all GTEx tissues (pooled medians) using Wilcoxon signed rank test in each tissue. As a result, we obtained *P*-values adjusted for testing in multiple tissues using the Benjamini-Hochberg (BH) correction and the sign of the deviation of each of the three metrics. Tissues, in which *e*_*l*_ and *e*_*g*_ had opposite signs, were discarded. For each USE, we selected tissues with significant (FDR < 0.05) and substantial (larger than 90% percentile in the absolute value) deviations of *ψ* and *e*_*g*_ and categorized them into two groups, tissues with opposite-sign deviations (expected for USE) and tissues with same-sign deviations (unexpected for USE). USEs with opposite-sign deviations in at least one tissue and no same-sign deviations in any tissue were considered tissue-specific. The list of GTEx tissues, their code designations, and colors is provided in Table S3.

### Identification of regulators in tissue-specific USEs

A number of large-scale functional assays have assessed the response of the cellular transcriptome to perturbations of RBP expression levels [41]. We collected a panel of 419 such experiments followed by RNA-seq for 248 RBPs (Table S4). Splicing responses to RBP expression perturbations were assessed by rMATS software separately for each experiment [48]. For uniformity, Δ*ψ* values reflecting the difference between the high and the low RBP condition (Control vs. KD, Control vs. KO, and OE vs. Control) were used. Only *ψ* values with *e*_*l*_ ≥ 15 and |Δ*ψ*| ≥ 0.1 were considered. Consequently, each USE was characterized by Δ*ψ* and Δ*e*_*g*_ values, one for each perturbation experiment, with their respective *P*-values after BH correction. To account for a different number of perturbation experiments for different RBPs, Δ*ψ* and Δ*e*_*g*_ values were converted into scores ±1.5, ±1, and 0, where the sign corresponds to the sign of Δ*ψ* and Δ*e*_*g*_, and the absolute values 1.5, 1, and 0 correspond to significant differences (FDR < 0.05), insignificant differences (FDR≥ 0.05), and discarded values, respectively. These scores were averaged over all perturbation experiments for each RBP-USE pair. An RBP was recognized as a candidate regulator of an USE if the average Δ*ψ* and Δ*e*_*g*_ scores had opposite signs. For each RBP, we also assessed *e*_*r*_, its expression level in GTEx tissues. To test whether tissue-specific RBP expression correlates with tissue-specific unproductive splicing, we used the same strategy as in the previous section but replacing *e*_*g*_ with *e*_*r*_.

### RBP footprinting data

The complete POSTAR3 database in tabular format was kindly provided by the authors [40]. An RBP-USE pair was considered as having CLIP support in the gene if the RBP had at least one peak obtained by Piranha or eCLIP pipelines within the gene that harbors the USE. An RBP-USE pair was further considered as having local CLIP support if a Piranha or eCLIP peak occurred within the intronic regions flanking the USE that were extended by 20 nts into the exons.

### Proteomic data

MS1-level data on protein expression in human tissues were downloaded from the Proteomics DB portal [52]. To test for negative associations between *ψ* and the expression of protein gene product, we matched tissues from Proteomics DB to GTEx tissues (Table S5), computed median *ψ* values for Proteomics DB tissues, and sorted tissues in ascending order by the median *ψ*. One-sided Jonckheere trend test was applied to check whether protein expression level follows a descending trend. Rejection of the null hypothesis in such a test indicated a negative association between *ψ* and the protein expression level.

### Genetic variation data

The GTEx portal provides lists of splicing quantitative trait loci (sQTLs) and expression quantitative trait loci (eQTLs) for within-tissue splicing and gene expression changes based on the genotype data from whole genome sequencing (WGS) [39]. Since sQTLs were produced with LeafCutter [53] that quantifies AS events based on split reads, we excluded intron retention and intron detention events from this analysis. Lists of sQTLs and eQTLs were mapped to USEs to identify splicing-and-expression quantitative trait loci (seQTLs), in which the minor allele is characterized by opposite changes of *ψ* and *e*_*g*_.

### Cell culture and siRNA transfection

Human A549 lung adenocarcinoma cells were cultured in Dulbecco’s modified Eagle’s medium/ Nutrient Mixture F-12 supplemented with 10% fetal bovine serum and 1% GlutaMAX (all products from Thermo Fisher Scientific). 1.5 ∗ 10^5^ cells were seeded in a 12-well plate for RNA extraction, and 3 ∗ 10^5^ cells were seeded in a 6-well plate for western blotting. Cells were transfected with 100 nM of a control siRNA targeting the firefly luciferase gene or with 100 nM of *PTBP1* targeting siRNA (Table S6) [54] for 24 h using Lipofectamine RNAiMAX transfection reagent (Invitrogen). Three hours before harvest, cycloheximide (CHX) was added to the cells giving a final concentration in the growth medium 300 µg/ml.

### RT-PCR

Total RNA was isolated by a guanidinium thiocyanate phenol-chloroform method using ExtractRNA Reagent (Evrogene) [55]. 1000 ng of total RNA was first subjected to RNase-Free DNase I digestion (Thermo Fisher Scientific) at 37^◦^C for 30 minutes to remove contaminating gDNA. Next, 500 ng of total RNA was used for cDNA synthesis Maxima First Strand cDNA Synthesis Kit for RT-qPCR (Thermo Fisher Scientific) to a total volume of 20 µl. cDNA was diluted 1:10 with nuclease-free water for qPCR analysis. qPCR reactions were performed in triplicates in a final volume of 12 µl in 96 well plates with 420 nM gene-specific primers, 2 µl of cDNA using Maxima SYBR Green/ROX qPCR Master Mix (2X) (Thermo Fisher Scientific). A sample without reverse transcriptase enzyme was included as a control to verify the absence of genomic DNA contamination. Amplification of the targets was carried out on CFX96 Real-Time system (Bio-Rad); cycle parameters were as follows: 95^◦^C for 2 min, followed by 45 cycles at 95^◦^C for 61^◦^C for 30 s, and 72^◦^C for 30 s, and ended at 72^◦^C for 5 min. A list of all primers used for qRT-PCR can be found in Table S6. The expression of genes and gene isoforms was calculated using assay-specific PCR efficiency. The result of the replicate measurement was considered an outlier and rejected if its *C*_*q*_ value was not in the range of 0.5 cycles [56].

Endpoint PCR was performed using 0.25 U recombinant Taq DNA Polymerase (Thermo Fisher Scientific) with 2 mM MgCl_2_, 1×KCl buffer, 2 µl of cDNA (same as for RT-qPCR), mM of each dNTPs, and 420 nM of each forward and reverse primer (Table S6). Cycling conditions were as follows: 95^◦^C for 5 min, followed by 35 cycles at 95^◦^C for 30 s, 61^◦^C for 30 s, and 72^◦^C for 30 s and ended at 72^◦^C for 5 min. PCR products were visualized on the agarose gel.

### Western blotting

Cell lysates were subjected to SDS-PAGE, and proteins were transferred to nitrocellulose membranes (Bio-Rad). Membranes were blocked in Tris-buffered saline (TBS, pH 7.4) containing 5% BSA, washed three times in TBS and incubated with primary anti-Ptbp1 antibody (Cloud-Clone Corp., PAC737Hu01) overnight at 4^◦^C (1:2000 dilution) followed by secondary Goat anti-Rabbit IgG (H+L) antibody HRP conjugate (Thermo Fisher Scientific, G21234) for 1h at room temperature (1:2000 dilution).

### Statistical analysis

The data were analyzed using python version 3.8.2 and R statistics software version 3.6.3. Non-parametric tests were performed with the scipy.stats python package using normal approximation with continuity correction. Jonckheere trend test was performed with the clinfun R package. In all figures, the significance levels of 5%, 1%, and 0.1% are denoted by *, **, and ***, respectively.

## RESULTS

To obtain a stringent list of USEs with tissue-specific regulation, we developed a six-step pipeline, at each successive step of which the set of USEs was gradually narrowed down (Figure 1). It starts with the extraction of candidate USEs from transcripts annotated as NMD targets according to the GENCODE database and literature search. In the second step, we identify USEs with significant negative association between NMD isoform splicing rate measured by the Percent-Spliced-In (PSI, *ψ*) metric and the expression level of the host gene (*e*_*g*_). These USEs are referred to as significant. In the third step, we identify tissues with significant deviations of *ψ* and *e*_*g*_ from the respective medians over all tissues. Significant opposite sign deviationsM Qare expected in unproductive splicing, while the same sign deviations represent a pattern that is unexpected. We classify USE as tissue-specific if it has a significant opposite sign deviation in at least one tissue and no significant same sign deviations. In the fourth step, we identify USEs that respond to RBP perturbations and check for the concordance of tissue-specific RBP expression levels and NMD isoform splicing rate in the fifth step. This yields a list of USEs, which we term tissue-specifically regulated. In the last step, we require evidence of binding for the predicted RBP-USE pairs in the host gene (CLIP support in the gene) and, more stringently, in the vicinity of USEs (local CLIP support) using the POSTAR3 database [40]. The former and the latter USEs are referred to as having CLIP support in the gene and having local CLIP support, respectively. The detailed description of each step is found in Methods. The number of USEs that remain after each step is shown in Table 1.

**Table 1:** The number of USEs.

| USE group | Validated | Annotated | Total |
| --- | --- | --- | --- |
| All | 48 | 2,754 | 2,802 |
| Significant | 11 | 568 | 579 |
| Tissue-specific | 5 | 132 | 137 |
| Tissue-specifically regulated | 4 | 80 | 84 |
| CLIP in the gene | 4 | 46 | 50 |
| Local CLIP support | 3 | 24 | 27 |

**Figure 1:**
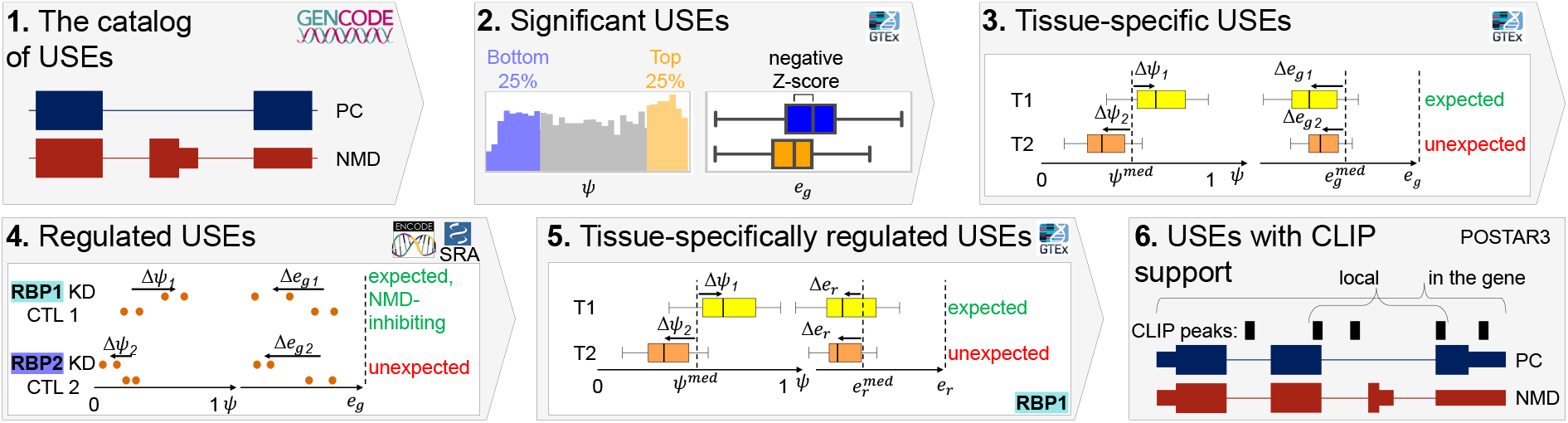
The workflow. The pipeline is designed to identify USEs and categorize them based on their splicing and expression patterns and evidence of regulation by RBPs. (1) An initial catalog of USEs is extracted from the transcriptome annotation. (2) Significant USEs are selected based on the association between *ψ* and *e*_*g*_. (3) Opposite sign deviations of *ψ* and *e*_*g*_ are expected in tissue-specific USEs. (4) Opposite sign deviations of *ψ* and *e*_*g*_ are expected in RBP perturbation experiments. The mode of regulation (NMD-inhibiting or NMD-promoting) is predicted for each RBP-USE pair. (5) Concordant changes of *ψ* and *e*_*r*_ are expected in tissue-specifically regulated USEs (6) Selection of USEs with CLIP support in the gene and local CLIP support.

### The USE catalog

In performing a literature search to catalog experimentally validated USEs, we collected information on the experimental outcomes and indicated whether an USE is reactive to NMD inactivation, whether RBP perturbations affect the NMD isoform inclusion or gene expression on protein and mRNA levels, and whether there is evidence of binding of the targeted mRNA by RBPs (Table S1). As a result, we obtained 237 RBP-USE pairs containing 48 RBPs and 57 USEs, including 203 cross-regulatory and 34 auto-regulatory cases, for which the experimental validation was reported in human or mouse cell lines or tissues. These USEs are referred to as validated; they are shown in Figures S4 and S5, where the latter illustrates a particularly dense subnetwork for SR proteins.

We considered four major types of AS events, namely cassette (skipped) exons, alternative 5’- and 3’-splice sites, and retained introns. Among them, we identified AS events leading to the generation of the NMD isoform (i.e., potential USEs) based on the GENCODE annotation [43] and classified them according to poison-essential dichotomy (Figure S1A). An USE is referred to as poison (essential) if the NMD isoform represents an insertion (deletion) to the protein-coding isoform. This definition extends the classification that was originally introduced for cassette exons to other types of AS events [57]. For instance, poison exons are normally skipped, but they trigger degradation by NMD when included in the mRNA. Numerous poison exons have been described in genes encoding SR proteins [9]. The reciprocal case is essential exons, which are normally included in the mRNA, but induce a PTC when skipped. Examples of essential exons are found in genes encoding hnRNP proteins, including PTBP1 [17], PTBP2 [18], and FUS [19]. Similarly, an alternative splice site or intron is referred to as poison (essential) if PTC occurs as a result of an insertion to (deletion from) the protein-coding isoform. Examples of these USEs have also been described [58, 57, 30, 59]. In total, 5,271 USEs, which will be referred to as annotated, were extracted from the GENCODE database (Table S2).

### Significant USEs

To study the association between *ψ* and e_*g*_, we analyzed RNA-seq data from about 8.5 thousand GTEx samples, first disregarding their tissue attribution. Samples from testis were excluded since the action of the NMD system in this tissue is different from that in other tissues [60, 61, 62]. After the removal of USEs with low variability and low read support, 2,754 out of 5,271 annotated cases and 48 out of 57 validated USEs remained (Table 1). To detect concordant changes between *ψ* and *e*_*g*_, we characterized each USE by *ψ*_*H*_−*ψ*_*L*_, the difference of the median *ψ* values, Δ*e*_*g*_, the difference of the median *e*_*g*_ values between the upper and the lower quartiles of the *ψ* distribution, and the corresponding *z*-score (see Methods). The comparison of metrics that distinguish USEs from non-NMD AS events revealed a reasonable threshold *z* < −5 yielding 579 significant USEs, in which an increase in *ψ* is accompanied by a significant drop in *e*_*g*_ (Table S7). Among them, there were 11 validated cases, most of which showed a remarkable magnitude of *ψ* changes (Figure 2A).

**Figure 2:**
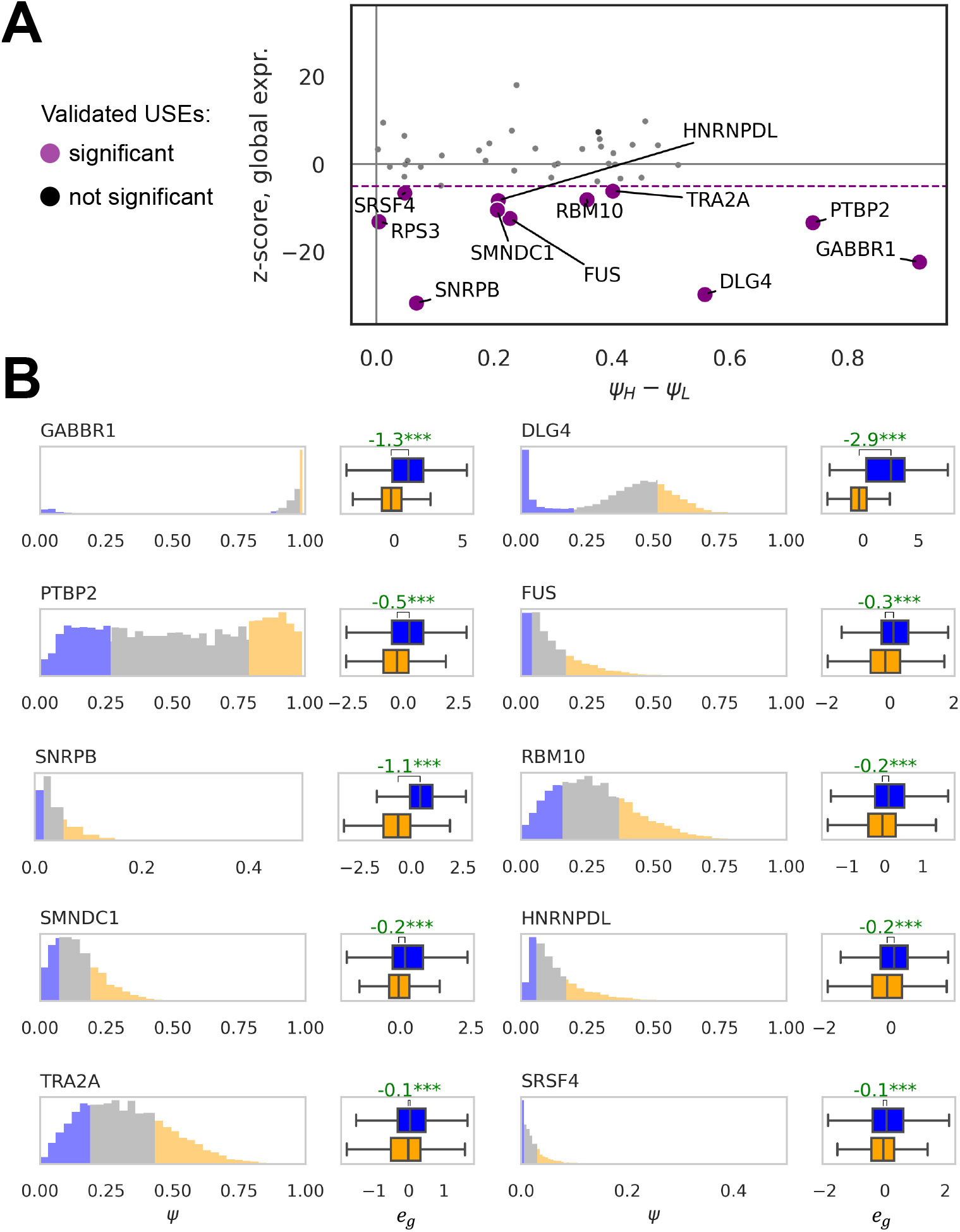
Significant validated USEs. **(A)** Significant USEs are characterized by the *z* < −5. *ψ*_*H*_ − *ψ*_*L*_ denotes the difference of the medians in the upper and the lower quartiles of the *ψ* distribution. **(B)** The distribution of *ψ* values for selected USEs from panel (A). The top 25% and the bottom 25% of *ψ* values are colored in orange and blue, respectively. Box plots represent the distribution of *e*_*g*_ in the two groups. Shown in green are Δ*e*_*g*_ values. Asterisks (***) denote statistically discernible differences at the 0.1% significance level.

The strongest association between *ψ* and *e*_*g*_ was observed for the well-documented *PTBP1* targets such as *GABBR1* and *DLG4*, in which *ψ*_*H*_ − *ψ*_*L*_ exceeded 50% and was accompanied by more than a twofold decrease of the expression level (Figure 2B). Other examples include USEs in *RBM10*, an important component of the spliceosome that is associated with genetic diseases and cancer [63, 64, 65], and *TRA2A*, a member of the SR protein family also known as an oncogene [66, 67, 68]. In *RBM10*, skipping of the essential exon 6 leads to a decrease in expression, which promotes proliferation in lung adenocarcinoma [69, 21]. In *TRA2A*, the suppression of expression occurs through the stimulation of the poison exon 2 in its mRNA by the *TRA2A* gene product and its paralog TRA2B [16]. These examples demonstrate that our analysis recapitulates unproductive splicing patterns in validated USEs. Remarkably, among annotated USEs we detected 16 cases, in which e_g_ change was as large as that in *GABBR1* and *DLG4* (Table S7).

### Tissue-specific regulation of USEs

To explore tissue-specificity of the association between *ψ* and *e*_*g*_ for significant USEs, we detected significant opposite-sign deviations of tissue medians of *ψ* and *e*_*g*_ from the pooled medians (see Methods). This analysis yielded 137 tissue-specific USEs (Table 1). Remarkably, cerebellum was characterized by simultaneous upregulation of NMD isoforms and downregulation of expression in many genes, while in brain, blood, muscle, and heart, the pattern was the contrary (Figure S6). This finding suggests a special role of NMD in orchestrating cerebellar transcriptional programs [70] amid generally higher abundance of AS events and their contribution to the development of all these tissues [71, 72, 73].

Next, a panel of large-scale functional assays was used to assess the concordance between *ψ* and *e*_*g*_ changes in response to RBP expression level perturbations [41]. We analyzed 248 RBPs (Table S4) and selected RBP-USE pairs, in which *ψ* and *e*_*g*_ changes were significant and had opposite signs (see Methods). This assessment was possible in 124 out of 203 validated cross-regulatory cases, among which we identified 30 RBP-USE pairs with the expected regulatory outcome and only one pair in which the direction of the regulation was opposite to that reported in the literature (Table S9). In application to tissue-specific USEs, it yielded at least one regulator in 113 cases, 84 of which were tissue-specifically regulated, i.e., tissue-specific RBP expression level (*e*_*r*_) was accompanied by tissue-specific *ψ* changes (Table 1).

We next asked whether RBP-USE networks are influenced by single nucleotide variants (SNVs) that potentially activate or suppress NMD isoforms. To address this, we identified splicing-and-expression quantitative trait loci (seQTLs), i.e., SNVs that significantly impact both splicing and gene expression in opposite directions (see Methods). Minor allele frequency (MAF) of seQTLs affecting tissue-specifically regulated USEs was substantially lower compared to that of seQTLs affecting other USEs (Figure S7B) indicating that SNVs could be disruptive for functionally important, tissue-specifically regulated USEs.

Since association doesn’t imply physical regulation, we further confined the list of candidates by requiring that RBPs bind the target pre-mRNA. Crosslinking and immunoprecipitation (CLIP) experiments within POSTAR3 database [40] presented evidence of binding by at least one regulatory RBP for 50 tissue-specifically regulated USEs, including 27 cases with footprints immediately proximal to USEs (Table 1, Table S10). In general, the predicted RBP-USE regulatory pairs were significantly enriched with RBP footprints both in the USE host gene and in the introns flanking the USE (Figure S8). The catalog of tissue-specifically regulated USEs with local CLIP support is available in Supplementary Data File 1.

### USE regulatory network

Clustering of 27 tissue-specifically regulated USEs with local CLIP support by their *ψ* values (Figure 3A) revealed four clusters that are characterized by decreased *ψ* in the brain (cluster 1), increased *ψ* in the brain (cluster 2), decreased *ψ* in blood (cluster 3), and decreased *ψ* in skeletal muscle and heart (cluster 4). These USEs belong to a regulatory network, in which *PTBP1* controls the expression of seven different targets (Figure 3B). Among them there are USEs in *DLG4* and *GABBR1* genes, which are suppressed in the brain and activated in other tissues [18, 38, 36]. In both these cases, we reconfirm that a brain-specific decrease in *ψ* and *e*_*r*_ is accompanied by an increase in *e*_*g*_, with a particularly strong trend on the protein level in the case of *DLG4*, which is consistent with the model of *PTBP1*-mediated skipping of essential exons (Figure S9).

**Figure 3:**
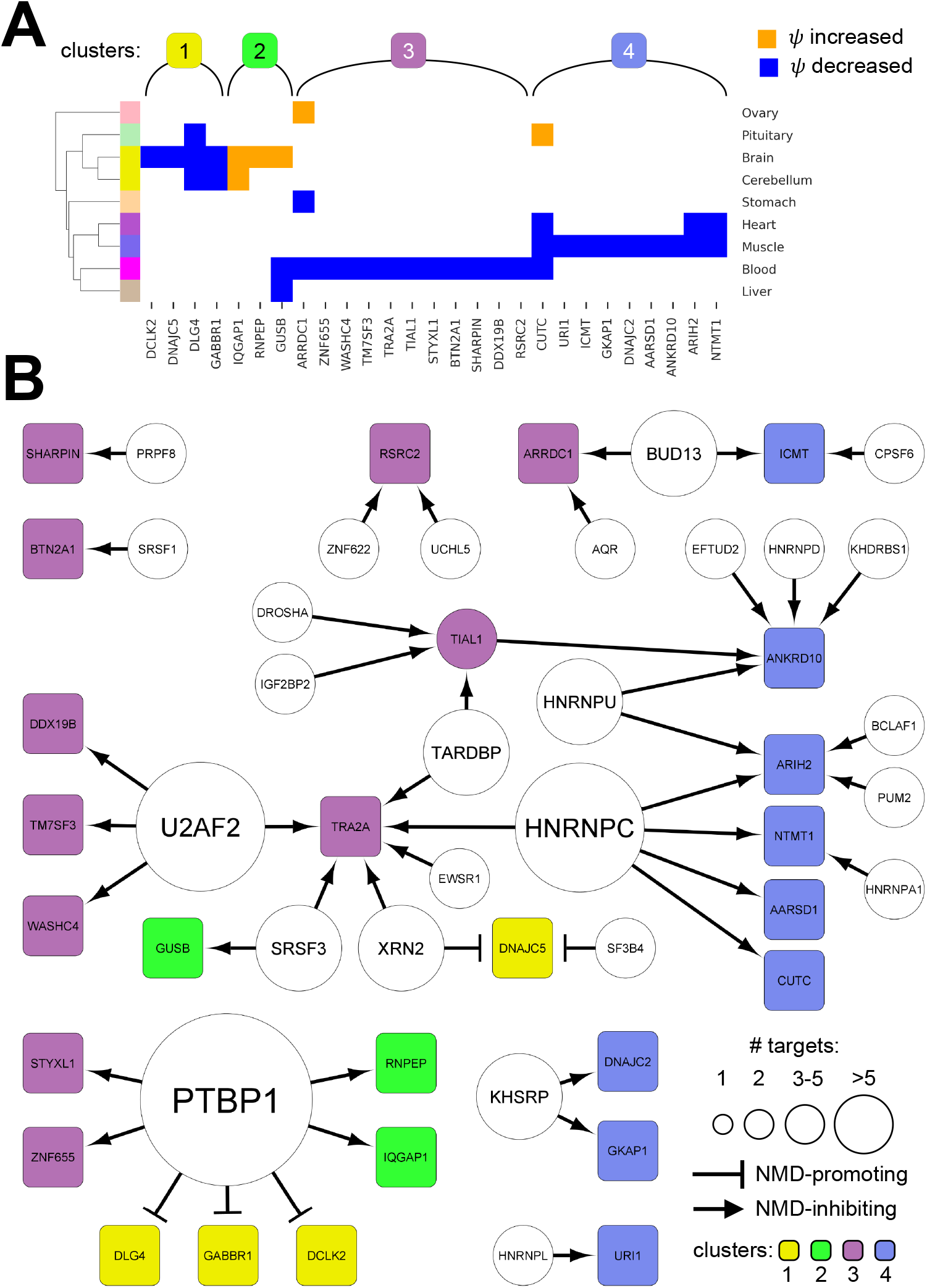
Tissue-specifically regulated USEs. **(A)** A clustering diagram of 27 tissue-specifically regulated USEs with local CLIP support. The clusters are characterized by decreased *ψ* in the brain (cluster 1), increased *ψ* in the brain (cluster 2), decreased *ψ* in blood (cluster 3), and decreased *ψ* in skeletal muscle and heart (cluster 4). **B** The predicted network of tissue-specifically regulated USEs with local CLIP support. The nodes represent USEs listed in panel (A). The edges represent NMD-promoting and NMD-inhibiting regulatory connections

We validated two examples of brain-specific USEs that are *PTBP1* targets (Figure 4). The first gene, *DCLK2*, is necessary for the development of the hippocampus and the regulation of dendrite remodeling [74, 75]. Gene expression and splicing patterns indicate that *PTBP1* stimulates the inclusion of the poison exon 16 in non-neural tissues, which leads to the suppression of *DCLK2* expression (Figure 4A, left, Figure S10, top). Together with the presence of CLIP peaks both upstream and downstream of the poison exon, this model assumes a non-canonical role of *PTBP1* as a splicing activator, consistently with the reports on the dual action of *PTBP1* that depends on its binding position relative to the exon [76]. To confirm the proposed regulatory model, we used siRNA-mediated knockdown (KD) of *PTBP1* in the A549 cell line and confirmed a stable ∼10-fold decrease in *PTBP1* expression using qRT-PCR and western blot analysis (Figure S11). We observed a substantial (∼ 25%) decrease in exon 16 inclusion rate upon siRNA treatment, both in the condition when NMD pathway was intact and inactivated by cycloheximide (CHX) (Figure 4A, right).

**Figure 4:**
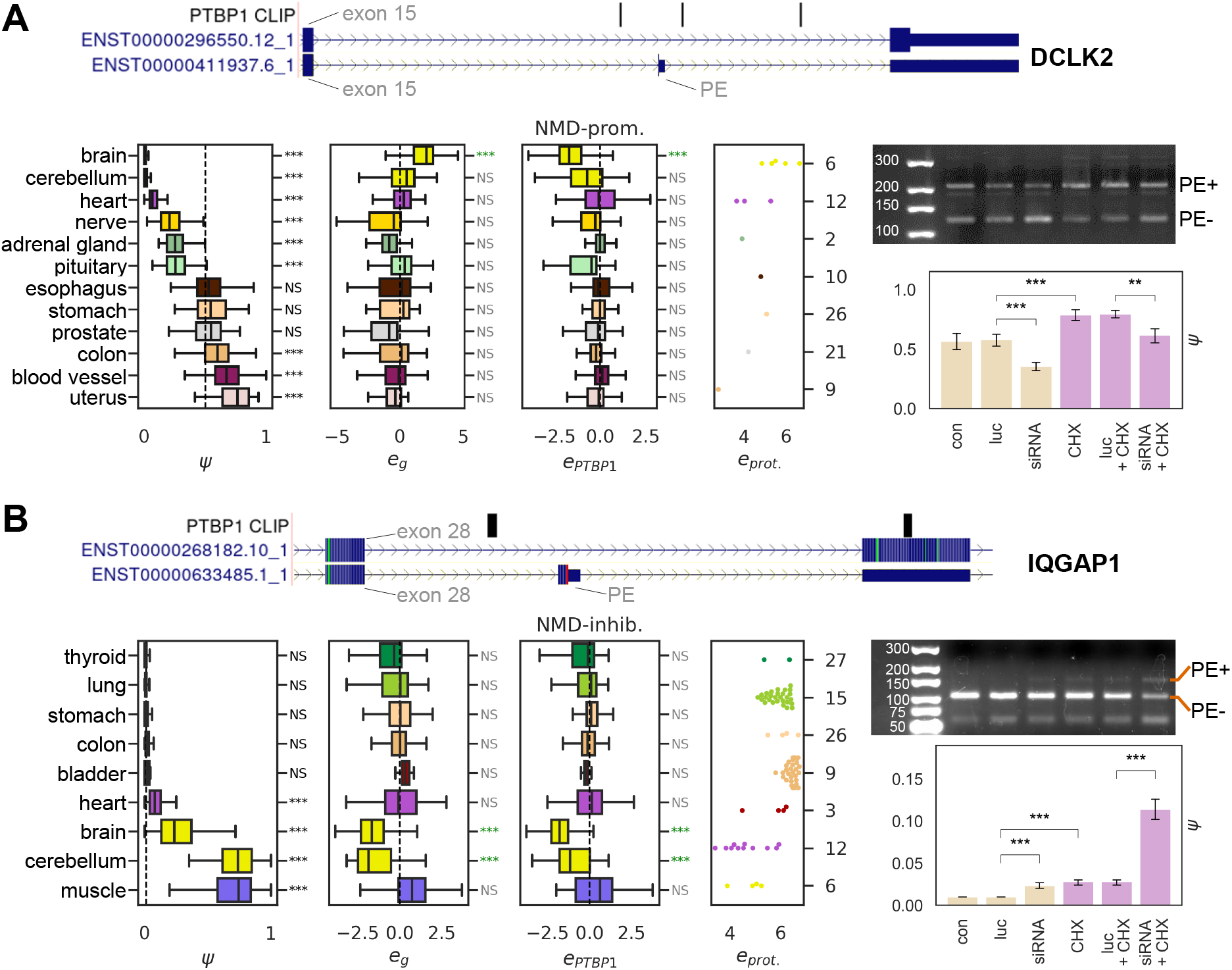
Examples of novel tissue-specific USEs. Panels (A) and (B) show the results for *DCLK2* and *IQGAP1* genes, respectively. In each case, boxplots show (left to right) the distribution of *ψ*, gene expression level (*e*_*g*_), the expression levels of the regulator (*PTBP1*), and the protein expression level. Statistically significant deviation from the pooled median are marked with asterisks (^∗∗∗^FDR < 0.001; ^∗∗^FDR < ^∗^FDR < 0.05; NS not significant). The color of an asterisk shows whether the deviation is in the expected (green) direction. Tissues are colored by GTEx color codes (Table S3) and sorted in ascending *ψ* order. The ideograms in each panel show the USE and CLIP peaks of *PTBP1*. The subpanels on the right show the results of RT-PCR (top) and qRT-PCR (bottom) experiments. “PE+” and “PE-” denote AS isoforms with and without the poison exon, respectively. The lanes are (left to right) non-treated control (con), treatment with siRNA targeting the firefly luciferase gene (luc), treatment with siRNA targeting *PTBP1*, CHX treatment, CHX and luc treatment, treatment with CHX and siRNA targeting *PTBP1*. The code for asterisks is as in the left subpanels.

The second validated gene, *IQGAP1*, is maintained at low levels in healthy brain tissues, but in brain tumors it is overexpressed [77]. We propose a model of brain-specific downregulation of *IQGAP1* by *PTBP1* through the inclusion of poison exon 29, which is almost completely suppressed in non-neural tissues (Figure 4B, left, Figure S10, bottom). The pattern of CLIP signal upstream of the USE suggests that *PTBP1*, which is downregulated in the brain, is a suppressor of exon 29 inclusion. In full agreement with this prediction, we observed an increase in exon 29 inclusion upon *PTBP1* KD, with a much stronger response during NMD inactivation by CHX (Figure 4, right). This observation points to a potential limitation of the proposed method, which has to do with selection bias against NMD isoforms that are actively degraded in tissues.

To further investigate into this bias, we used publicly available RNA-seq data on siRNA-mediated PTBP1 KD with fractionation into cytoplasmic and nuclear RNA in Hela cells [47] to monitor the predicted targets of *PTBP1*. As expected, statistically significant splicing changes were observed exclusively in the nuclear fraction, while statistically significant gene expression changes were observed exclusively in the cytoplasmic fraction (Figure S12). Remarkably, the magnitude of differential expression was considerably larger in the cytoplasmic fraction, e.g., *GABBR1* was five times more abundant in the cytoplasm than in the nuclear fraction. The concordance between cytoplasmic and nuclear responses, with the exception of *DCLK2* which, however, was not significant, demonstrates overall applicability of the method.

## DISCUSS1ON

A coordinated interaction between AS and NMD that leads to the degradation of specific mRNAs, which constitutes unproductive splicing, is a widespread phenomenon that has been documented in almost all eukaryotes [78, 79, 11, 15]. Compared to other post-transcriptional regulatory mechanisms such as endogenous RNA interference [80, 81, 82, 83] and the control of mRNA stability by RNA modifications [84, 85, 86], unproductive splicing appears to act pervasively at the transcriptome level, as evidenced by the fact that nearly a third of human protein-coding genes have at least one annotated NMD transcript isoform, and many of these isoforms exhibit evolutionary conservation [87].

A fundamental challenge in studying USEs is the existence of feedback loops and indirect connections in splicing networks. In autoregulation, for instance, the product of a gene that harbors a poison exon counteracts its own upregulation by promoting exon inclusion, thus leading to the downregulation of gene expression level and, consequently, to suppression of the NMD isoform that was initially upregulated. Splicing regulatory networks may contain circuits that serve as proxies and interfere with direct interactions, e.g., *PTBP1* upregulates *SRSF3*, which in turn upregulates *PTBP2*, but *PTBP1* itself directly downregulates *PTBP2* (Figure S4). Interestingly, RBP perturbation experiments consistently demonstrated nearly the same number of positive and negative associations between *ψ* and *e*_*g*_ in the validated autoregulatory USEs (Table S9) implying the existence of positive feedback loops, the physiological relevance of which is debatable [57]. Consequently, the sign and the magnitude of the association between NMD isoform splicing rate and RBP expression may vary depending on the connectivity in the network potentially leading to both false positive and false negative predictions.

Significant associations between *ψ* and *e*_*g*_ were observed in many USE, however the majority of validated USEs have no evidence of such association. In fact, unproductive splicing can often be implicit on the RNA level, as several experimental studies failed to detect changes in mRNA expression despite substantial changes in protein abundance (examples in Table S1). In part, this lack of response could be explained by a bias in observing splicing changes that are masked by the degradation of unproductive isoforms by the NMD pathway and constant influx of pre-mRNAs due to ongoing transcription [27]. Unproductive splicing could also be inactive in fully differentiated tissues or operate only in specific conditions such as differentiation [88, 12], neurogenesis [38, 18], or hypoxia [89]. In fact, most splicing factors demonstrate only minor differences in expression between tissues [90] in comparison to KD or OE experiments, in which the validated USEs have been originally described. Consequently, the regulatory potential of unproductive splicing requires a relatively large magnitude of AS changes, which may be restricted in mature tissues. All these factors inherently limit the sensitivity of the proposed method.

The methodology outlined here consists of six steps, which sequentially check for associations between AS and host gene expression, AS responses to RBP perturbations, associations between AS and RBP expression, and RBP footprints near USE. It relies on large amounts of RNA-seq data and implements multifactorial, stringent filtration. The method yielded a stringent list of 27 tissue-specifically regulated USEs with CLIP support, which likely represents only a tip of the unproductive splicing iceberg, and two USEs from this list were confirmed experimentally. Further analysis of molecular mechanisms underlying unproductive splicing and increasing volumes of multi-omics data will likely expand the unproductive splicing network outlined in this study.

## CONCLUSION

This study presents a bioinformatic workflow for the identification of tissue-specific USEs and their regulators. It confirms the regulatory mechanisms in validated cases and predicts 27 novel regulatory USEs including *PTBP1* targets in the *DCLK2* and *IQGAP1*, for which we provide experimental validation. These results greatly expand the current knowledge on tissue-specific regulation of unproductive splicing and open new avenues for future research.

## AVAILABILITY OF DATA AND MATERIALS

The datasets generated during the current study are available online at https://doi.org/10.5281/zenodo.6786643. The source code used for the analysis is available at https://github.com/magmir71/unproductive_splicing.

## ACKNOWLEDGEMENTS

The authors thank Drs. Roderic Guigo’, Silvia Perez Lluch, and Juan Valca’rcel for insightful discussions about molecular mechanisms of unproductive splicing. A.M., M.V., S.M., A.A.M., and D.D.P. acknowledge Russian Science Foundation grant 22-14-00330.

## References

[1] Lykke-Andersen, S. and Jensen, T. H. (Nov, 2015) Nonsense-mediated mRNA decay: an intricate machinery that shapes transcriptomes. Nat Rev Mol Cell Biol, 16(11), 665–677.

[2] Medghalchi, S. M., Frischmeyer, P. A., Mendell, J. T., Kelly, A. G., Lawler, A. M., and Dietz, H. C. (Jan, 2001) Rent1, a trans-effector of nonsense-mediated mRNA decay, is essential for mammalian embryonic viability. Hum Mol Genet, 10(2), 99–105.

[3] Weischenfeldt, J., Damgaard, ı., Bryder, D., Theilgaard-Mönch, K., Thoren, L. A., Nielsen, F. C., Jacobsen, S. E., Nerlov, C., and Porse, B. T. (May, 2008) NMD is essential for hematopoietic stem and progenitor cells and for eliminating by-products of programmed DNA rearrangements. Genes Dev, 22(10), 1381–1396.

[4] Mcılwain, D. R., Pan, Q., Reilly, P. T., Elia, A. J., McCracken, S., Wakeham, A. C., ıtie-Youten, A., Blencowe, B. J., and Mak, T. W. (Jul, 2010) Smg1 is required for embryogenesis and regulates diverse genes via alternative splicing coupled to nonsense-mediated mRNA decay. Proc Natl Acad Sci U S A, 107(27), 12186–12191.

[5] Li, T., Shi, Y., Wang, P., Guachalla, L. M., Sun, B., Joerss, T., Chen, Y. S., Groth, M., Krueger, A., Platzer, M., Yang, Y. G., Rudolph, K. L., and Wang, Z. Q. (Jun, 2015) Smg6/Est1 licenses embryonic stem cell differentiation via nonsense-mediated mRNA decay. EMBO J, 34(12), 1630–1647.

[6] Lou, C. H., Shao, A., Shum, E. Y., Espinoza, J. L., Huang, L., Karam, R., and Wilkinson, M. F. (Feb, 2014) Posttranscriptional control of the stem cell and neurogenic programs by the nonsense-mediated RNA decay pathway. Cell Rep, 6(4), 748–764.

[7] Lewis, B. P., Green, R. E., and Brenner, S. E. (Jan, 2003) Evidence for the widespread coupling of alternative splicing and nonsense-mediated mRNA decay in humans. Proc Natl Acad Sci U S A, 100(1), 189–192.

[8] Desai, A., Hu, Z., French, C. E., Lloyd, J. P. B., and Brenner, S. E. (2020) Networks of Splice Factor Regulation by Unproductive Splicing Coupled With Nonsense Mediated mRNA Decay. bioRxiv,.

[9] Lareau, L. F., ınada, M., Green, R. E., Wengrod, J. C., and Brenner, S. E. (Apr, 2007) Unproductive splicing of SR genes associated with highly conserved and ultraconserved DNA elements. Nature, 446(7138), 926–929.

[10] Filichkin, S. A. and Mockler, T. C. (Jul, 2012) Unproductive alternative splicing and nonsense mRNAs: a widespread phenomenon among plant circadian clock genes. Biol Direct, 7, 20.

[11] Ni, J. Z., Grate, L., Donohue, J. P., Preston, C., Nobida, N., O’Brien, G., Shiue, L., Clark, T. A., Blume, J. E., and Ares, M. (Mar, 2007) Ultraconserved elements are associated with homeostatic control of splicing regulators by alternative splicing and nonsense-mediated decay. Genes Dev, 21(6), 708–718.

[12] Wong, J. J., Ritchie, W., Ebner, O. A., Selbach, M., Wong, J. W., Huang, Y., Gao, D., Pinello, N., Gonzalez, M., Baidya, K., Thoeng, A., Khoo, T. L., Bailey, C. G., Holst, J., and Rasko, J. E. (Aug, 2013) Orchestrated intron retention regulates normal granulocyte differentiation. Cell, 154(3), 583–595.

[13] Dewaele, M., Tabaglio, T., Willekens, K., Bezzi, M., Teo, S. X., Low, D. H., Koh, C. M., Rambow, F., Fiers, M., Rogiers, A., Radaelli, E., Al-Haddawi, M., Tan, S. Y., Hermans, E., Amant, F., Yan, H., Lakshmanan, M., Koumar, R. C., Lim, S. T., Derheimer, F. A., Campbell, R. M., Bonday, Z., Tergaonkar, V., Shackleton, M., Blattner, C., Marine, J. C., and Guccione, E. (Jan, 2016) Antisense oligonucleotide-mediated MDM4 exon 6 skipping impairs tumor growth. J Clin ınvest, 126(1), 68–84.

[14] Barbier, J., Dutertre, M., Bittencourt, D., Sanchez, G., Gratadou, L., de la Grange, P., and Auboeuf, D. (Oct, 2007) Regulation of H-ras splice variant expression by cross talk between the p53 and nonsense-mediated mRNA decay pathways. Mol Cell Biol, 27(20), 7315–7333.

[15] Garc’ıa-Moreno, J. F. and Romaõ, L. (Dec, 2020) Perspective in Alternative Splicing Coupled to Nonsense-Mediated mRNA Decay. ınt J Mol Sci, 21(24).

[16] Leclair, N. K., Brugiolo, M., Urbanski, L., Lawson, S. C., Thakar, K., Yurieva, M., George, J., Hinson, J. T., Cheng, A., Graveley, B. R., and Anczuko’w, O. (11, 2020) Poison Exon Splicing Regulates a Coordinated Network of SR Protein Expression during Differentiation and Tumorigenesis. Mol Cell, 80(4), 648–665.

[17] Spellman, R., Llorian, M., and Smith, C. W. (Aug, 2007) Crossregulation and functional redundancy between the splicing regulator PTB and its paralogs nPTB and ROD1. Mol Cell, 27(3), 420–434.

[18] Boutz, P. L., Stoilov, P., Li, Q., Lin, C. H., Chawla, G., Ostrow, K., Shiue, L., Ares, M., and Black, D. L. (Jul, 2007) A post-transcriptional regulatory switch in polypyrimidine tract-binding proteins reprograms alternative splicing in developing neurons. Genes Dev, 21(13), 1636–1652.

[19] Zhou, Y., Liu, S., Liu, G., Oztürk, A., and Hicks, G. G. (Oct, 2013) ALS-associated FUS mutations result in compromised FUS alternative splicing and autoregulation. PLoS Genet, 9(10), e1003895.

[20] Kemmerer, K., Fischer, S., and Weigand, J. E. (03, 2018) Auto- and cross-regulation of the hnRNPs D and DL. RNA, 24(3), 324–331.

[21] Sun, Y., Bao, Y., Han, W., Song, F., Shen, X., Zhao, J., Zuo, J., Saffen, D., Chen, W., Wang, Z., You, X., and Wang, Y. (Aug, 2017) Autoregulation of RBM10 and cross-regulation of RBM10/RBM5 via alternative splicing-coupled nonsense-mediated decay. Nucleic Acids Res, 45(14), 8524–8540.

[22] Saltzman, A. L., Pan, Q., and Blencowe, B. J. (Feb, 2011) Regulation of alternative splicing by the core spliceosomal machinery. Genes Dev, 25(4), 373–384.

[23] Cuccurese, M., Russo, G., Russo, A., and Pietropaolo, C. (2005) Alternative splicing and nonsense-mediated mRNA decay regulate mammalian ribosomal gene expression. Nucleic Acids Res, 33(18), 5965–5977.

[24] Takei, S., Togo-Ohno, M., Suzuki, Y., and Kuroyanagi, H. (07, 2016) Evolutionarily conserved autoregulation of alternative pre-mRNA splicing by ribosomal protein L10a. Nucleic Acids Res, 44(12), 5585–5596.

[25] Lareau, L. F. and Brenner, S. E. (Apr, 2015) Regulation of splicing factors by alternative splicing and NMD is conserved between kingdoms yet evolutionarily flexible. Mol Biol Evol, 32(4), 1072–1079.

[26] Kalyna, M., Simpson, C. G., Syed, N. H., Lewandowska, D., Marquez, Y., Kusenda, B., Marshall, J., Fuller, J., Cardle, L., McNicol, J., Dinh, H. Q., Barta, A., and Brown, J. W. (Mar, 2012) Alternative splicing and nonsense-mediated decay modulate expression of important regulatory genes in Arabidopsis. Nucleic Acids Res, 40(6), 2454–2469.

[27] Kovalak, C., Donovan, S., Bicknell, A. A., Metkar, M., and Moore, M. J. (05, 2021) Deep sequencing of pre-translational mRNPs reveals hidden flux through evolutionarily conserved alternative splicing nonsense-mediated decay pathways. Genome Biol, 22(1), 132.

[28] Wollerton, M. C., Gooding, C., Wagner, E. J., Garcia-Blanco, M. A., and Smith, C. W. (Jan, 2004) Autoregulation of polypyrimidine tract binding protein by alternative splicing leading to nonsense-mediated decay. Mol Cell, 13(1), 91–100.

[29] Jumaa, H. and Nielsen, P. J. (Aug, 1997) The splicing factor SRp20 modifies splicing of its own mRNA and ASF/SF2 antagonizes this regulation. EMBO J, 16(16), 5077–5085.

[30] Änkö, M. L., Müller-McNicoll, M., Brandl, H., Curk, T., Gorup, C., Henry, ı., Ule, J., and Neugebauer, K. M. (2012) The RNA-binding landscapes of two SR proteins reveal unique functions and binding to diverse RNA classes. Genome Biol, 13(3), R17.

[31] Jangi, M., Boutz, P. L., Paul, P., and Sharp, P. A. (Mar, 2014) Rbfox2 controls autoregulation in RNA-binding protein networks. Genes Dev, 28(6), 637–651.

[32] Rossbach, O., Hung, L. H., Schreiner, S., Grishina, ı., Heiner, M., Hui, J., and Bindereif, A. (Mar, 2009) Auto- and cross-regulation of the hnRNP L proteins by alternative splicing. Mol Cell Biol, 29(6), 1442–1451.

[33] Sun, S., Zhang, Z., Sinha, R., Karni, R., and Krainer, A. R. (Mar, 2010) SF2/ASF autoregulation involves multiple layers of post-transcriptional and translational control. Nat Struct Mol Biol, 17(3), 306–312.

[34] Jime’nez, M., Urtasun, R., Elizalde, M., Azkona, M., Latasa, M. U., Uriarte, ı., Arechederra, M., Alignani, D., Ba’rcena-Varela, M., A’lvarez Sola, G., Colyn, L., Santamar’ıa, E., Sangro, B., Rodriguez-Ortigosa, C., Ferna’ndez-Barrena, M. G., A’vila, M. A., and Berasain, C. (04, 2019) Splicing events in the control of genome integrity: role of SLU7 and truncated SRSF3 proteins. Nucleic Acids Res, 47(7), 3450–3466.

[35] Nakano, Y., Kelly, M. C., Rehman, A. U., Boger, E. T., Morell, R. J., Kelley, M. W., Friedman, T. B., and Ba’nfi, B. (07, 2018) Defects in the Alternative Splicing-Dependent Regulation of REST Cause Deafness. Cell, 174(3), 536–548.

[36] Zheng, S., Gray, E. E., Chawla, G., Porse, B. T., O’Dell, T. J., and Black, D. L. (Jan, 2012) PSD-95 is post-transcriptionally repressed during early neural development by PTBP1 and PTBP2. Nat Neurosci, 15(3), 381–388.

[37] Zheng, S. (Dec, 2016) Alternative splicing and nonsense-mediated mRNA decay enforce neural specific gene expression. ınt J Dev Neurosci, 55, 102–108.

[38] Makeyev, E. V., Zhang, J., Carrasco, M. A., and Maniatis, T. (Aug, 2007) The MicroRNA miR-124 promotes neuronal differentiation by triggering brain-specific alternative pre-mRNA splicing. Mol Cell, 27(3), 435–448.

[39] Aguet, F., Anand, S., Ardlie, K. G., Gabriel, S., Getz, G. A., Graubert, A., Hadley, K., Handsaker, R. E., Huang, K. H., Kashin, S., Li, X., MacArthur, D. G., Meier, S. R., Nedzel, J. L., Nguyen, D. T., Segrè, A. V., Todres, E., Balliu, B., Barbeira, A. N., Battle, A., Bonazzola, R., Brown, A., Brown, C. D., Castel, S. E., Conrad, D. F., Cotter, D. J., Cox, N., Das, S., de Goede, O. M., Dermitzakis, E. T., Einson, J., Engelhardt, B. E., Eskin, E., Eulalio, T. Y., Ferraro, N. M., Flynn, E. D., Fresard, L., Gamazon, E. R., Garrido-Mart’ın, D., Gay, N. R., Gloudemans, M. J., Guigo’, R., Hame, A. R., He, Y., Hoffman, P. J., Hormozdiari, F., Hou, L., ım, H. K., Jo, B., Kasela, S., Kellis, M., Kim-Hellmuth, S., Kwong, A., Lappalainen, T., Li, X., Liang, Y., Mangul, S., Mohammadi, P., Montgomery, S. B., Munõz-Aguirre, M., Nachun, D. C., Nobel, A. B., Oliva, M., Park, Y., Park, Y., Parsana, P., Rao, A. S., Reverter, F., Rouhana, J. M., Sabatti, C., Saha, A., Stephens, M., Stranger, B. E., Strober, B. J., Teran, N. A., Viñuela, A., Wang, G., Wen, X., Wright, F., Wucher, V., Zou, Y., Ferreira, P. G., Li, G., Mele’, M., Yeger-Lotem, E., Barcus, M. E., Bradbury, D., Krubit, T., McLean, J. A., Qi, L., Robinson, K., Roche, N. V., Smith, A. M., Sobin, L., Tabor, D. E., Undale, A., Bridge, J., Brigham, L. E., Foster, B. A., Gillard, B. M., Hasz, R., Hunter, M., Johns, C., Johnson, M., Karasik, E., Kopen, G., Leinweber, N W. F., McDonald, A., Moser, M. T., Myer, K., Ramsey, K. D., Roe, B., Shad, S., Thomas, J. A., Walters, G., Washington, M., Wheeler, J., Jewell, S. D., Rohrer, D. C., Valley, D. R., Davis, D. A., Mash, D. C., Branton, P. A., Barker, L. K., Gardiner, H. M., Mosavel, M., Siminoff, L. A., Flicek, P., Haeussler, M., Juettemann, T., Kent, W. J., Lee, C. M., Powell, C. C., Rosenbloom, K. R., Ruffier, M., Sheppard, D., Taylor, K., Trevanion, S. J., Zerbino, D. R., Abell, N. S., Akey, J., Chen, L., Demanelis, K., Doherty, J. A., Feinberg, A. P., Hansen, K. D., Hickey, P. F., Jasmine, F., Jiang, L., Kaul, R., Kibriya, M. G., Li, J. B., Li, Q., Lin, S., Linder, S. E., Pierce, B. L., Rizzardi, L. F., Skol, A. D., Smith, K. S., Snyder, M., Stamatoyannopoulos, J., Tang, H., Wang, M., Carithers, L. J., Guan, P., Koester, S. E., Little, A. R., Moore, H. M., Nierras, C. R., Rao, A. K., Vaught, J. B., and Volpi, S. (09, 2020) The GTEx Consortium atlas of genetic regulatory effects across human tissues. Science, 369(6509), 1318–1330.

[40] Zhao, W., Zhang, S., Zhu, Y., Xi, X., Bao, P., Ma, Z., Kapral, T. H., Chen, S., Zagrovic, B., Yang, Y. T., and Lu, Z. J. (01, 2022) POSTAR3: an updated platform for exploring post-transcriptional regulation coordinated by RNA-binding proteins. Nucleic Acids Res, 50(D1), D287–D294.

[41] Van Nostrand, E. L., Freese, P., Pratt, G. A., Wang, X., Wei, X., Xiao, R., Blue, S. M., Chen, J. Y., Cody, N. A. L., Dominguez, D., Olson, S., Sundararaman, B., Zhan, L., Bazile, C., Bouvrette, L. P. B., Bergalet, J., Duff, M. O., Garcia, K. E., Gelboin-Burkhart, C., Hochman, M., Lambert, N. J., Li, H., McGurk, M. P., Nguyen, T. B., Palden, T., Rabano, ı., Sathe, S., Stanton, R., Su, A., Wang, R., Yee, B. A., Zhou, B., Louie, A. L., Aigner, S., Fu, X. D., Le’cuyer, E., Burge, C. B., Graveley, B. R., and Yeo, G. W. (07, 2020) A large-scale binding and functional map of human RNA-binding proteins. Nature, 583(7818), 711–719.

[42] Haeussler, M., Zweig, A. S., Tyner, C., Speir, M. L., Rosenbloom, K. R., Raney, B. J., Lee, C. M., Lee, B. T., Hinrichs, A. S., Gonzalez, J. N., Gibson, D., Diekhans, M., Clawson, H., Casper, J., Barber, G. P., Haussler, D., Kuhn, R. M., and Kent, W. J. (01, 2019) The UCSC Genome Browser database: 2019 update. Nucleic Acids Res, 47(D1), D853–D858.

[43] Frankish, A., Diekhans, M., Jungreis, ı., Lagarde, J., Loveland, J. E., Mudge, J. M., Sisu, C., Wright, J. C., Armstrong, J., Barnes, ı., Berry, A., Bignell, A., Boix, C., Carbonell Sala, S., Cunningham, F., Di Domenico, T., Donaldson, S., Fiddes, ı. T., Garc’ıa Giro’n, C., Gonzalez, J. M., Grego, T., Hardy, M., Hourlier, T., Howe, K. L., Hunt, T., ızuogu, O. G., Johnson, R., Martin, F. J., Mart’ınez, L., Mohanan, S., Muir, P., Navarro, F. C. P., Parker, A., Pei, B., Pozo, F., Riera, F. C., Ruffier, M., Schmitt, B. M., Stapleton, E., Suner, M. M., Sycheva, ı., Uszczynska-Ratajczak, B., Wolf, M. Y., Xu, J., Yang, Y. T., Yates, A., Zerbino, D., Zhang, Y., Choudhary, J. S., Gerstein, M., Guigo’, R., Hubbard, T. J. P., Kellis, M., Paten, B., Tress, M. L., and Flicek, P. (01, 2021) GENCODE 2021. Nucleic Acids Res, 49(D1), D916–D923.

[44] Dobin, A., Davis, C. A., Schlesinger, F., Drenkow, J., Zaleski, C., Jha, S., Batut, P., Chaisson, M., and Gingeras, T. R. (Jan, 2013) STAR: ultrafast universal RNA-seq aligner. Bioinformatics, 29(1), 15–21.

[45] Mele’, M., Ferreira, P. G., Reverter, F., DeLuca, D. S., Monlong, J., Sammeth, M., Young, T. R., Goldmann, J. M., Pervouchine, D. D., Sullivan, T. J., Johnson, R., Segrè, A. V., Djebali, S., Niarchou, A., Wright, F. A., Lappalainen, T., Calvo, M., Getz, G., Dermitzakis, E. T., Ardlie, K. G., and Guigo’, R. (May, 2015) Human genomics. The human transcriptome across tissues and individuals. Science, 348(6235), 660–665.

[46] Pervouchine, D. D., Knowles, D. G., and Guigo’, R. (Jan, 2013) ıntron-centric estimation of alternative splicing from RNA-seq data. Bioinformatics, 29(2), 273–274.

[47] Attig, J., Agostini, F., Gooding, C., Chakrabarti, A. M., Singh, A., Haberman, N., Zagalak, J. A., Emmett, W., Smith, C. W. J., Luscombe, N. M., and Ule, J. (08, 2018) Heteromeric RNP Assembly at LıNEs Controls Lineage-Specific RNA Processing. Cell, 174(5), 1067– 1081.

[48] Shen, S., Park, J. W., Lu, Z. X., Lin, L., Henry, M. D., Wu, Y. N., Zhou, Q., and Xing, Y. (Dec, 2014) rMATS: robust and flexible detection of differential alternative splicing from replicate RNA-Seq data. Proc Natl Acad Sci U S A, 111(51), E5593–5601.

[49] Love, M. ı., Huber, W., and Anders, S. (2014) Moderated estimation of fold change and dispersion for RNA-seq data with DESeq2. Genome Biol, 15(12), 550.

[50] Graubert, A., Aguet, F., Ravi, A., Ardlie, K. G., and Getz, G. (Mar, 2021) RNA-SeQC 2: Efficient RNA-seq quality control and quantification for large cohorts. Bioinformatics,.

[51] Zhu, A., ıbrahim, J. G., and Love, M. ı. (06, 2019) Heavy-tailed prior distributions for sequence count data: removing the noise and preserving large differences. Bioinformatics, 35(12), 2084–2092.

[52] Lautenbacher, L., Samaras, P., Muller, J., Grafberger, A., Shraideh, M., Rank, J., Fuchs, S. T., Schmidt, T. K., The, M., Dallago, C., Wittges, H., Rost, B., Krcmar, H., Kuster, B., and Wilhelm, M. (01, 2022) ProteomicsDB: toward a FAıR open-source resource for life-science research. Nucleic Acids Res, 50(D1), D1541–D1552.

[53] Li, Y. ı., Knowles, D. A., Humphrey, J., Barbeira, A. N., Dickinson, S. P., ım, H. K., and Pritchard, J. K. (01, 2018) Annotation-free quantification of RNA splicing using LeafCutter. Nat Genet, 50(1), 151–158.

[54] Cho, C. Y., Chung, S. Y., Lin, S., Huang, J. S., Chen, Y. L., Jiang, S. S., Cheng, L. C., Kuo, T. H., Lay, J. D., Yang, Y. Y., Lai, G. M., and Chuang, S. E. (11, 2019) PTBP1-mediated regulation of AXL mRNA stability plays a role in lung tumorigenesis. Sci Rep, 9(1), 16922.

[55] Chomczynski, P. and Sacchi, N. (Apr, 1987) Single-step method of RNA isolation by acid guanidinium thiocyanate-phenol-chloroform extraction. Anal Biochem, 162(1), 156–159.

[56] de Ronde, M. W. J., Ruijter, J. M., Lanfear, D., Bayes-Genis, A., Kok, M. G. M., Creemers, E. E., Pinto, Y. M., and Pinto-Sietsma, S. J. (05, 2017) Practical data handling pipeline improves performance of qPCR-based circulating miRNA measurements. RNA, 23(5), 811– 821.

[57] Pervouchine, D., Popov, Y., Berry, A., Borsari, B., Frankish, A., and Guigo’, R. (06, 2019) ıntegrative transcriptomic analysis suggests new autoregulatory splicing events coupled with nonsense-mediated mRNA decay. Nucleic Acids Res, 47(10), 5293–5306.

[58] Hamid, F. M. and Makeyev, E. V. (Nov, 2014) Regulation of mRNA abundance by polypyrimidine tract-binding protein-controlled alternate 5’ splice site choice. PLoS Genet, 10(11), e1004771.

[59] Boutz, P. L., Bhutkar, A., and Sharp, P. A. (Jan, 2015) Detained introns are a novel, widespread class of post-transcriptionally spliced introns. Genes Dev, 29(1), 63–80.

[60] Shum, E. Y., Jones, S. H., Shao, A., Dumdie, J., Krause, M. D., Chan, W. K., Lou, C. H., Espinoza, J. L., Song, H. W., Phan, M. H., Ramaiah, M., Huang, L., McCarrey, J. R., Peterson, K. J., De Rooij, D. G., Cook-Andersen, H., and Wilkinson, M. F. (Apr, 2016) The Antagonistic Gene Paralogs Upf3a and Upf3b Govern Nonsense-Mediated RNA Decay. Cell, 165(2), 382–395.

[61] MacDonald, C. C. and Grozdanov, P. N. (May, 2017) Nonsense in the testis: multiple roles for nonsense-mediated decay revealed in male reproduction. Biol Reprod, 96(5), 939–947.

[62] Zetoune, A. B., Fontanière, S., Magnin, D., Anczuko’w, O., Buisson, M., Zhang, C. X., and Mazoyer, S. (Dec, 2008) Comparison of nonsense-mediated mRNA decay efficiency in various murine tissues. BMC Genet, 9, 83.

[63] Hegele, A., Kamburov, A., Grossmann, A., Sourlis, C., Wowro, S., Weimann, M., Will, C. L., Pena, V., Lührmann, R., and Stelzl, U. (Feb, 2012) Dynamic protein-protein interaction wiring of the human spliceosome. Mol Cell, 45(4), 567–580.

[64] Johnston, J. J., Teer, J. K., Cherukuri, P. F., Hansen, N. F., Loftus, S. K., Chong, K., Mullikin, J. C., and Biesecker, L. G. (May, 2010) Massively parallel sequencing of exons on the X chromosome identifies RBM10 as the gene that causes a syndromic form of cleft palate. Am J Hum Genet, 86(5), 743–748.

[65] Jung, J. H., Lee, H., Cao, B., Liao, P., Zeng, S. X., and Lu, H. (01, 2020) RNA-binding motif protein 10 induces apoptosis and suppresses proliferation by activating p53. Oncogene, 39(5), 1031–1040.

[66] Zhao, X., Chen, Q., Cai, Y., Chen, D., Bei, M., Dong, H., and Xu, J. (2021) TRA2A Binds With LncRNA MALAT1 To Promote Esophageal Cancer Progression By Regulating EZH2/β-catenin Pathway. J Cancer, 12(16), 4883–4890.

[67] Tan, Y., Hu, X., Deng, Y., Yuan, P., Xie, Y., and Wang, J. (10, 2018) TRA2A promotes proliferation, migration, invasion and epithelial mesenchymal transition of glioma cells. Brain Res Bull, 143, 138–144.

[68] Xu, W., Huang, H., Yu, L., and Cao, L. (Apr, 2015) Meta-analysis of gene expression profiles indicates genes in spliceosome pathway are up-regulated in hepatocellular carcinoma (HCC). Med Oncol, 32(4), 96.

[69] Zhao, J., Sun, Y., Huang, Y., Song, F., Huang, Z., Bao, Y., Zuo, J., Saffen, D., Shao, Z., Liu, W., and Wang, Y. (01, 2017) Functional analysis reveals that RBM10 mutations contribute to lung adenocarcinoma pathogenesis by deregulating splicing. Sci Rep, 7, 40488.

[70] Kurosaki, T., Sakano, H., Pröschel, C., Wheeler, J., Hewko, A., and Maquat, L. E. (11, 2021) NMD abnormalities during brain development in the Fmr1-knockout mouse model of fragile X syndrome. Genome Biol, 22(1), 317.

[71] Xu, Q., Modrek, B., and Lee, C. (Sep, 2002) Genome-wide detection of tissue-specific alternative splicing in the human transcriptome. Nucleic Acids Res, 30(17), 3754–3766.

[72] Yeo, G., Holste, D., Kreiman, G., and Burge, C. B. (2004) Variation in alternative splicing across human tissues. Genome Biol, 5(10), R74.

[73] Baralle, F. E. and Giudice, J. (07, 2017) Alternative splicing as a regulator of development and tissue identity. Nat Rev Mol Cell Biol, 18(7), 437–451.

[74] Kerjan, G., Koizumi, H., Han, E. B., Dube’, C. M., Djakovic, S. N., Patrick, G. N., Baram, T. Z., Heinemann, S. F., and Gleeson, J. G. (Apr, 2009) Mice lacking doublecortin and doublecortin-like kinase 2 display altered hippocampal neuronal maturation and spontaneous seizures. Proc Natl Acad Sci U S A, 106(16), 6766–6771.

[75] Shin, E., Kashiwagi, Y., Kuriu, T., ıwasaki, H., Tanaka, T., Koizumi, H., Gleeson, J. G., and Okabe, S. (2013) Doublecortin-like kinase enhances dendritic remodelling and negatively regulates synapse maturation. Nat Commun, 4, 1440.

[76] Hamid, F. M. and Makeyev, E. V. (12, 2017) A mechanism underlying position-specific regulation of alternative splicing. Nucleic Acids Res, 45(21), 12455–12468.

[77] McDonald, K. L., O’Sullivan, M. G., Parkinson, J. F., Shaw, J. M., Payne, C. A., Brewer, J. M., Young, L., Reader, D. J., Wheeler, H. T., Cook, R. J., Biggs, M. T., Little, N. S., Teo, C., Stone, G., and Robinson, B. G. (May, 2007) ıQGAP1 and ıGFBP2: valuable biomarkers for determining prognosis in glioma patients. J Neuropathol Exp Neurol, 66(5), 405– 417.

[78] Mitrovich, Q. M. and Anderson, P. (Sep, 2000) Unproductively spliced ribosomal protein mRNAs are natural targets of mRNA surveillance in C. elegans. Genes Dev, 14(17), 2173– 2184.

[79] Hansen, K. D., Lareau, L. F., Blanchette, M., Green, R. E., Meng, Q., Rehwinkel, J., Gallusser, F. L., ızaurralde, E., Rio, D. C., Dudoit, S., and Brenner, S. E. (Jun, 2009) Genome-wide identification of alternative splice forms down-regulated by nonsense-mediated mRNA decay in Drosophila. PLoS Genet, 5(6), e1000525.

[80] Wight, M. and Werner, A. (2013) The functions of natural antisense transcripts. Essays Biochem, 54, 91–101.

[81] Guo, Z., Maki, M., Ding, R., Yang, Y., Zhang, B., and Xiong, L. (Jun, 2014) Genomewide survey of tissue-specific microRNA and transcription factor regulatory networks in 12 tissues. Sci Rep, 4, 5150.

[82] Watanabe, T., Totoki, Y., Toyoda, A., Kaneda, M., Kuramochi-Miyagawa, S., Obata, Y., Chiba, H., Kohara, Y., Kono, T., Nakano, T., Surani, M. A., Sakaki, Y., and Sasaki, H. (May, 2008) Endogenous siRNAs from naturally formed dsRNAs regulate transcripts in mouse oocytes. Nature, 453(7194), 539–543.

[83] Tam, O. H., Aravin, A. A., Stein, P., Girard, A., Murchison, E. P., Cheloufi, S., Hodges, E., Anger, M., Sachidanandam, R., Schultz, R. M., and Hannon, G. J. (May, 2008) Pseudogene-derived small interfering RNAs regulate gene expression in mouse oocytes. Nature, 453(7194), 534–538.

[84] Yu, J., Chen, M., Huang, H., Zhu, J., Song, H., Zhu, J., Park, J., and Ji, S. J. (02, 2018) Dynamic m6A modification regulates local translation of mRNA in axons. Nucleic Acids Res, 46(3), 1412–1423.

[85] Ke, S., Pandya-Jones, A., Saito, Y., Fak, J. J., Vågbø, C. B., Geula, S., Hanna, J. H., Black, D. L., Darnell, J. E., and Darnell, R. B. (05, 2017) A mRNA modifications are deposited in nascent pre-mRNA and are not required for splicing but do specify cytoplasmic turnover. Genes Dev, 31(10), 990–1006.

[86] Licht, K., Hartl, M., Amman, F., Anrather, D., Janisiw, M. P., and Jantsch, M. F. (01, 2019) ınosine induces context-dependent recoding and translational stalling. Nucleic Acids Res, 47(1), 3–14.

[87] Mudge, J. M., Frankish, A., and Harrow, J. (Dec, 2013) Functional transcriptomics in the post-ENCODE era. Genome Res, 23(12), 1961–1973.

[88] Kim, E., ılagan, J. O., Liang, Y., Daubner, G. M., Lee, S. C., Ramakrishnan, A., Li, Y., Chung, Y. R., Micol, J. B., Murphy, M. E., Cho, H., Kim, M. K., Zebari, A. S., Aumann, S., Park, C. Y., Buonamici, S., Smith, P. G., Deeg, H. J., Lobry, C., Aifantis, ı., Modis, Y., Allain, F. H., Halene, S., Bradley, R. K., and Abdel-Wahab, O. (May, 2015) SRSF2 Mutations Contribute to Myelodysplasia by Mutant-Specific Effects on Exon Recognition. Cancer Cell, 27(5), 617–630.

[89] Gardner, L. B. (Jun, 2008) Hypoxic inhibition of nonsense-mediated RNA decay regulates gene expression and the integrated stress response. Mol Cell Biol, 28(11), 3729–3741.

[90] Gerstberger, S., Hafner, M., Ascano, M., and Tuschl, T. (2014) Evolutionary conservation and expression of human RNA-binding proteins and their role in human genetic disease. Adv Exp Med Biol, 825, 1–55.

