## Supplementary Information for "Tissue-specific regulation of gene expression via unproductive splicing"

November 21, 2022

### List of Supplementary Data Files

SDF1. A gallery of 27 tissue-specifically regulated USEs with strong evidence of RBP binding.

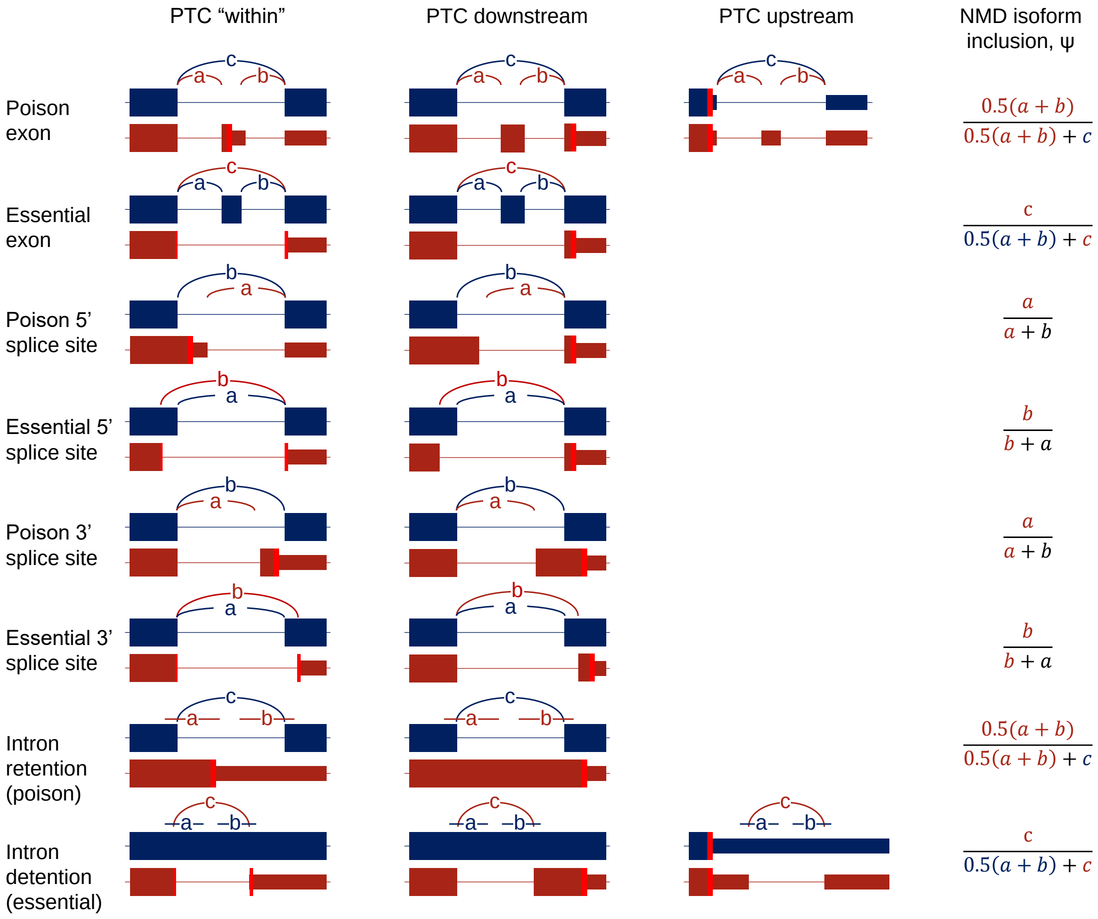

**Figure S1:** The four major AS types, cassette (skipped) exons, alternative 5’- and 3’-splice sites, and retained introns, give rise to poison and essential USEs. The NMD isoform is shown in dark red; the protein-coding isoform is shown in dark blue. Red vertical stripe denotes a PTC, which can be within the AS event (first column) or occur downstream as a result of a frameshift (second column). In rare cases, a PTC can also occur upstream of the AS event and be recognized as an NMD target due to a splice junction more than 50 nts downstream (third column). The definition of the $\psi$ value is shown in the fourth column for all USE classes; $a$, $b$, and $c$ denote the number of split reads.

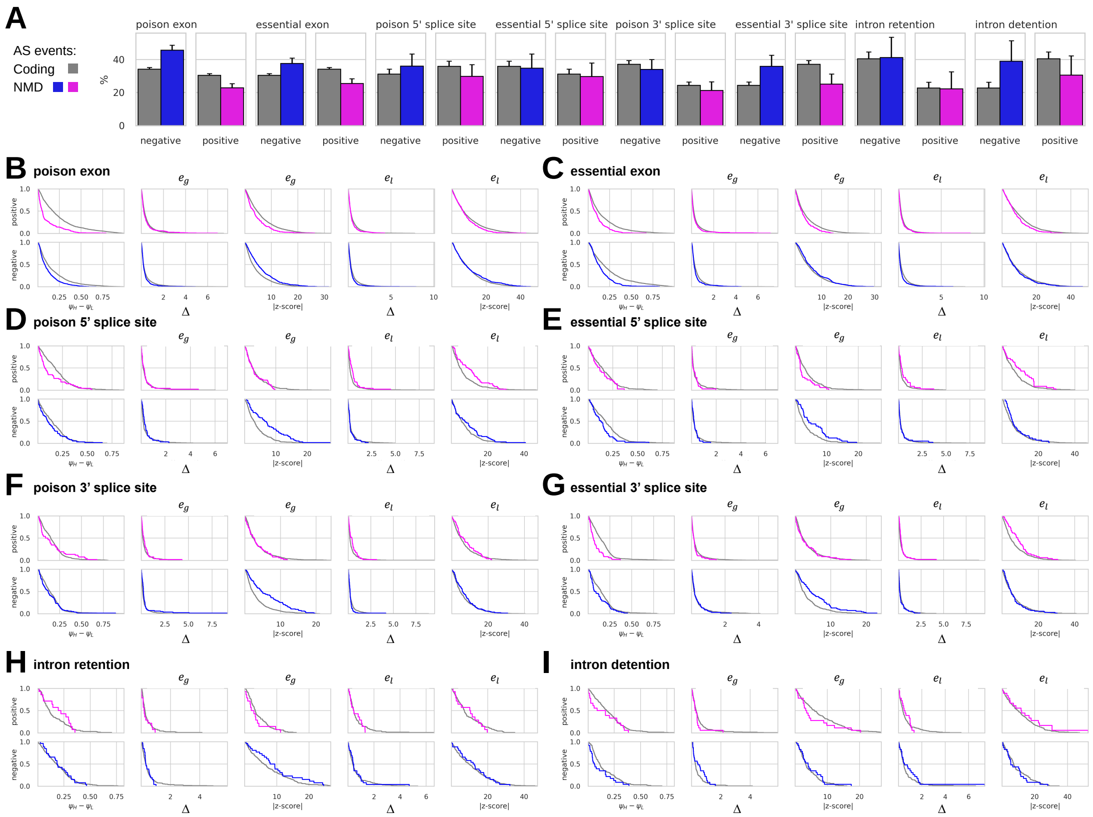

**Figure S2:** Discriminating features of USEs vs. protein-coding AS events. **A** The fraction of USEs and protein-coding AS events with the same sign of $\Delta e_g$ and $\Delta e_l$ (positive, $\Delta e_g > 0$ and $\Delta e_l > 0$, and negative, $\Delta e_g < 0$ and $\Delta e_l < 0$). Error bars represent 95% confidence intervals. **(B-I)** The distribution of $\psi_H - \psi_L$, $\Delta e_g$, $z - score$ of $e_g$, $\Delta e_l$, $z - score$ of $e_l$ in events from the positive and negative sets (see panel A). The empirical cumulative distribution function ($CDF$) is shown as $1 - CDF$.

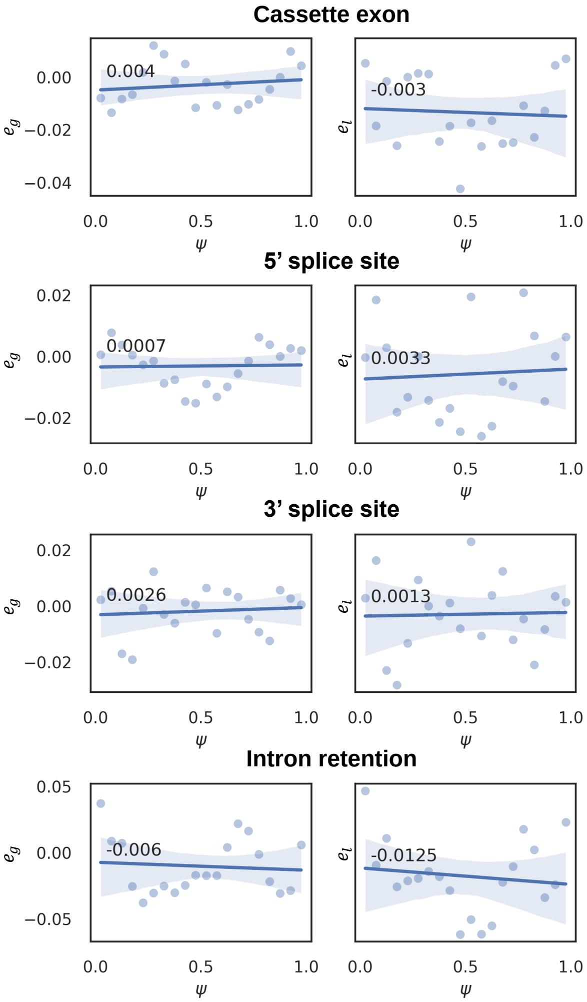

**Figure S3:** The dependence of the median $e_g$ and $e_l$ values on $\psi$ in protein-coding AS events. The $x$-axis represents $\psi$ values, which were subdivided into 20 bins of equal size. The $y$-axis represents median $e_g$ (left) and $e_l$ (right) values in $\psi$ bins. Optimal least square regression line (blue) and the 95% confidence interval (gray) are shown. None of the AS classes shows a significant dependence of median $e_g$ and $e_l$ values on $\psi$.

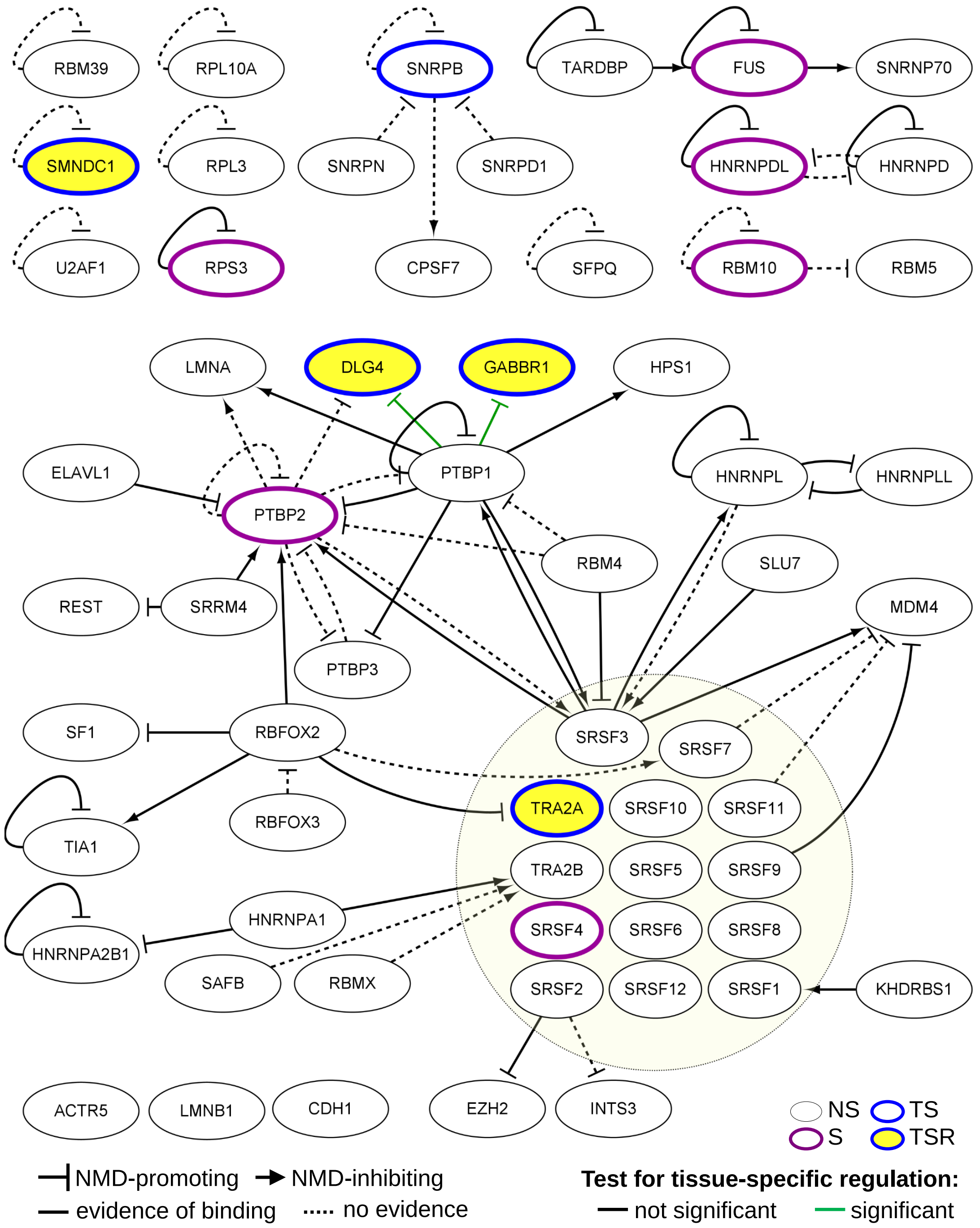

**Figure S4:** The network of validated USEs. Nodes represent USEs listed in Table S1. Edges represent NMD-promoting and NMD-inhibiting regulatory connections. The color code of the nodes represents the classification of USEs as significant (S), tissue-specific (TS), or tissue-specifically regulated (TSR) (Table 1). Edges are colored according to tissue-specificity. The subnetwork of SR proteins is exempt to Figure S5.

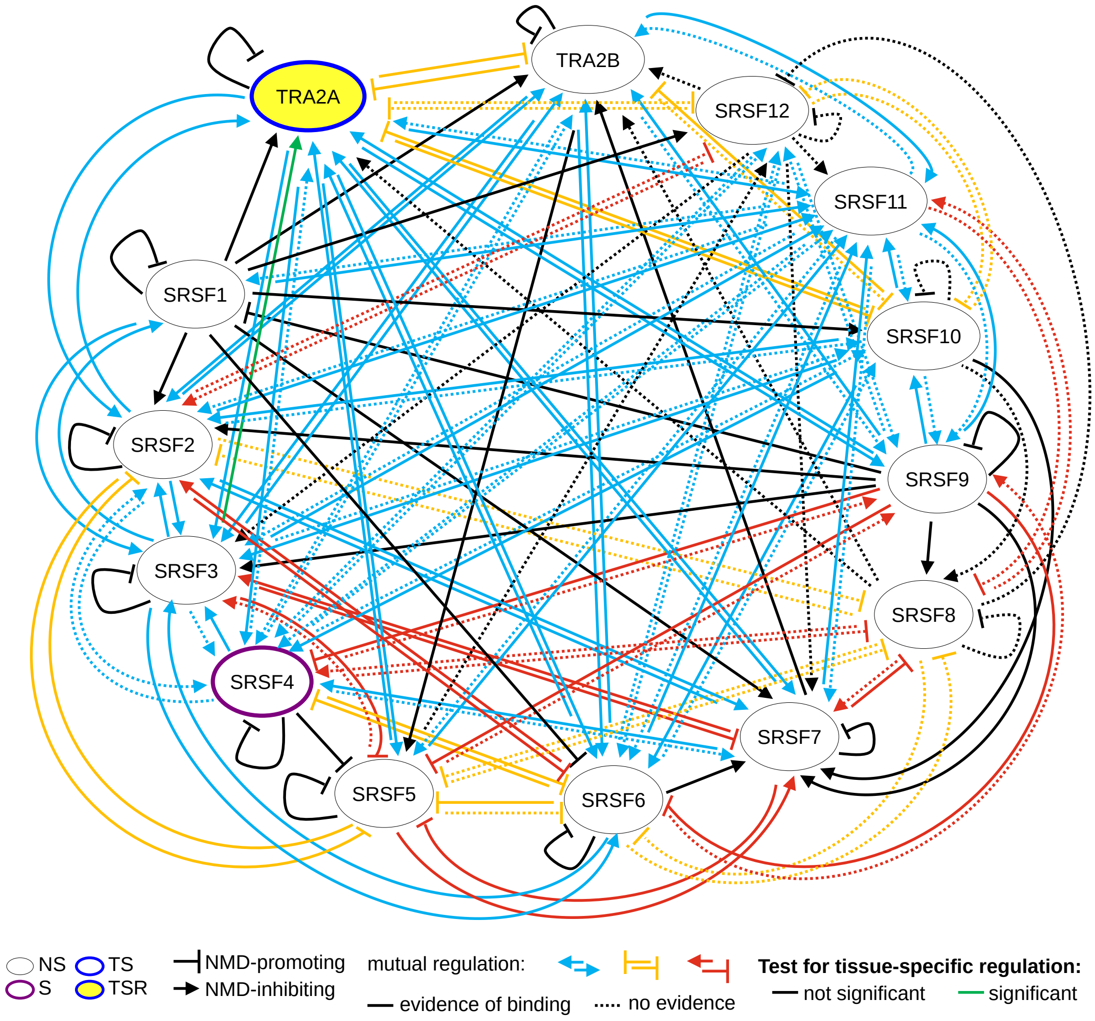

**Figure S5:** The regulatory subnetwork of SR proteins according to the data from Table S1. The legend is as in Figure S4.

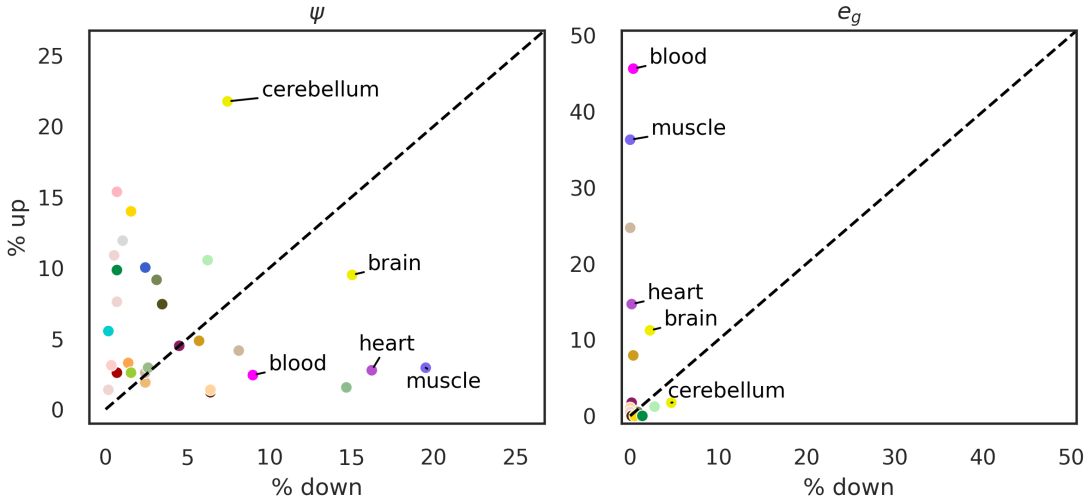

**Figure S6:** Characterization of tissues by the number of up- and downregulated USEs and genes. **(A)** The fraction of significant USEs that are upregulated ($y$-axis) and downregulated ($x$-axis). Tissue color codes are shown in Table S3. **(B)** The fraction of significant USEs in which the gene is upregulated ($y$-axis) or downregulated ($x$-axis).

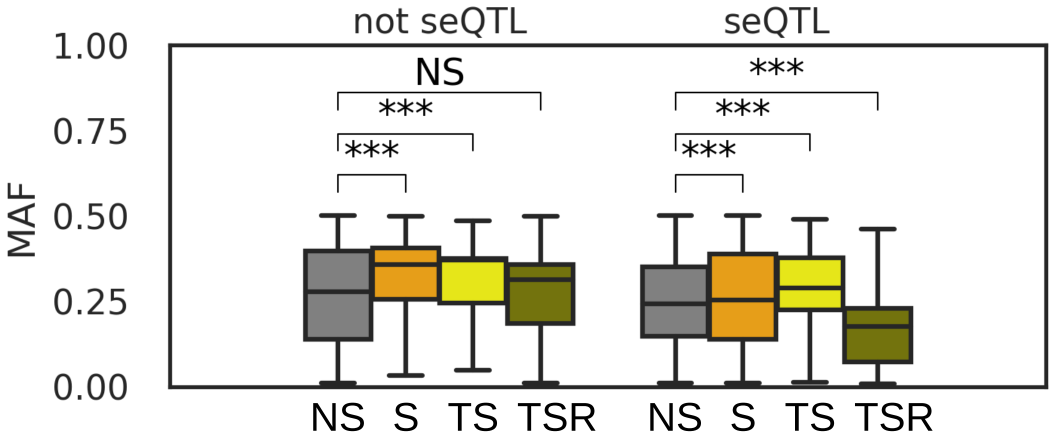

**Figure S7:** sQTLs that affect USE classes (Table 1). Minor allele frequency (MAF) of seQTLs (right) and non-seQTLs (left) associated in four USE classes: not significant (NS, gray); significant but not tissue-specific (S, orange); tissue-specific but not tissue-specifically regulated (TS, yellow); tissue-specifically regulated (TSR, olive). Asterisks denote statistical significance of the Mann-Whitney sum of ranks test (***P $< 0.001$; NS not significant).

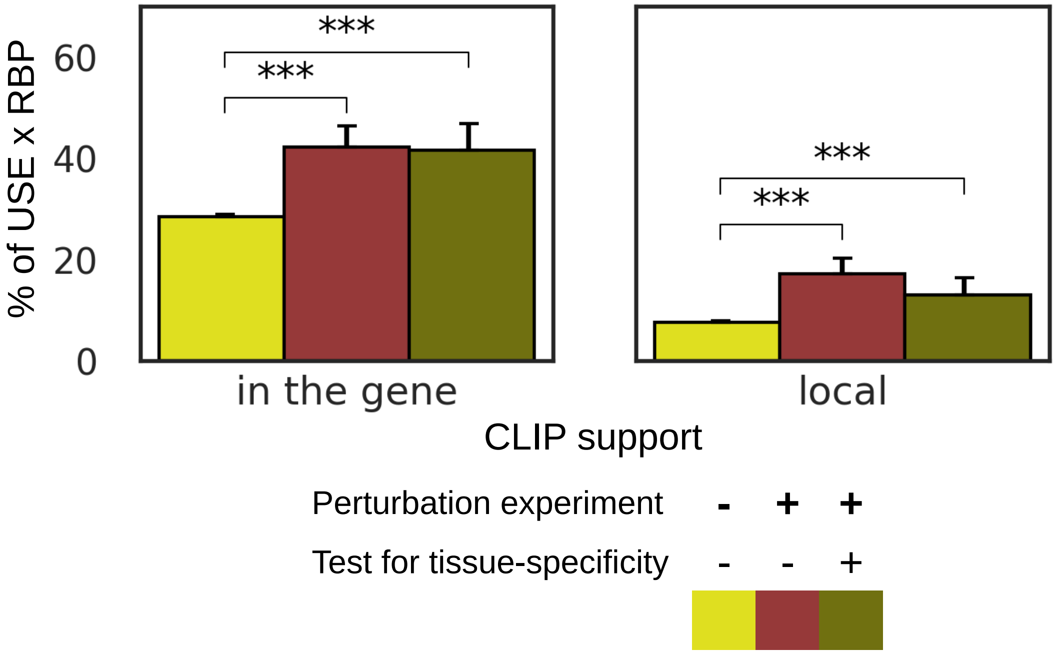

**Figure S8:** Presence of CLIP peaks and USE responses to RBP perturbations. All pairs of tissue-specific USEs and RBPs listed in Table S4 were classified by the response to RBP perturbation and tissue-specificity. The fractions of USE-RBP pairs with CLIP peak in the gene (left) and local CLIP peak (right) were compared using Fisher exact test (***P $< 0.001$; NS not significant).

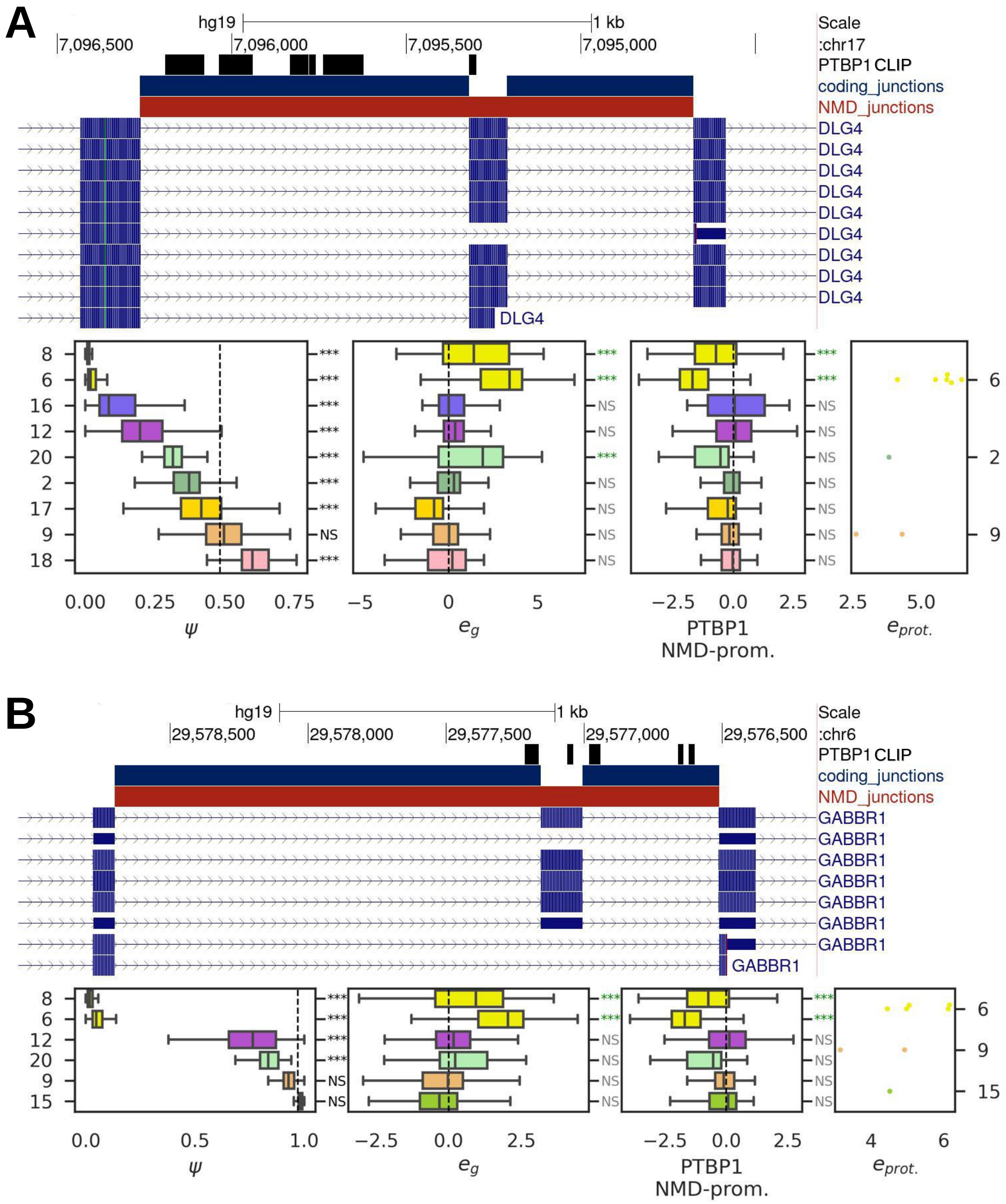

**Figure S9:** Examples of validated tissue-specific USEs. Panels (A) and (B) show the results for *DLG4* and *GABBR1* genes, respectively. The legend is as in Figure 4.

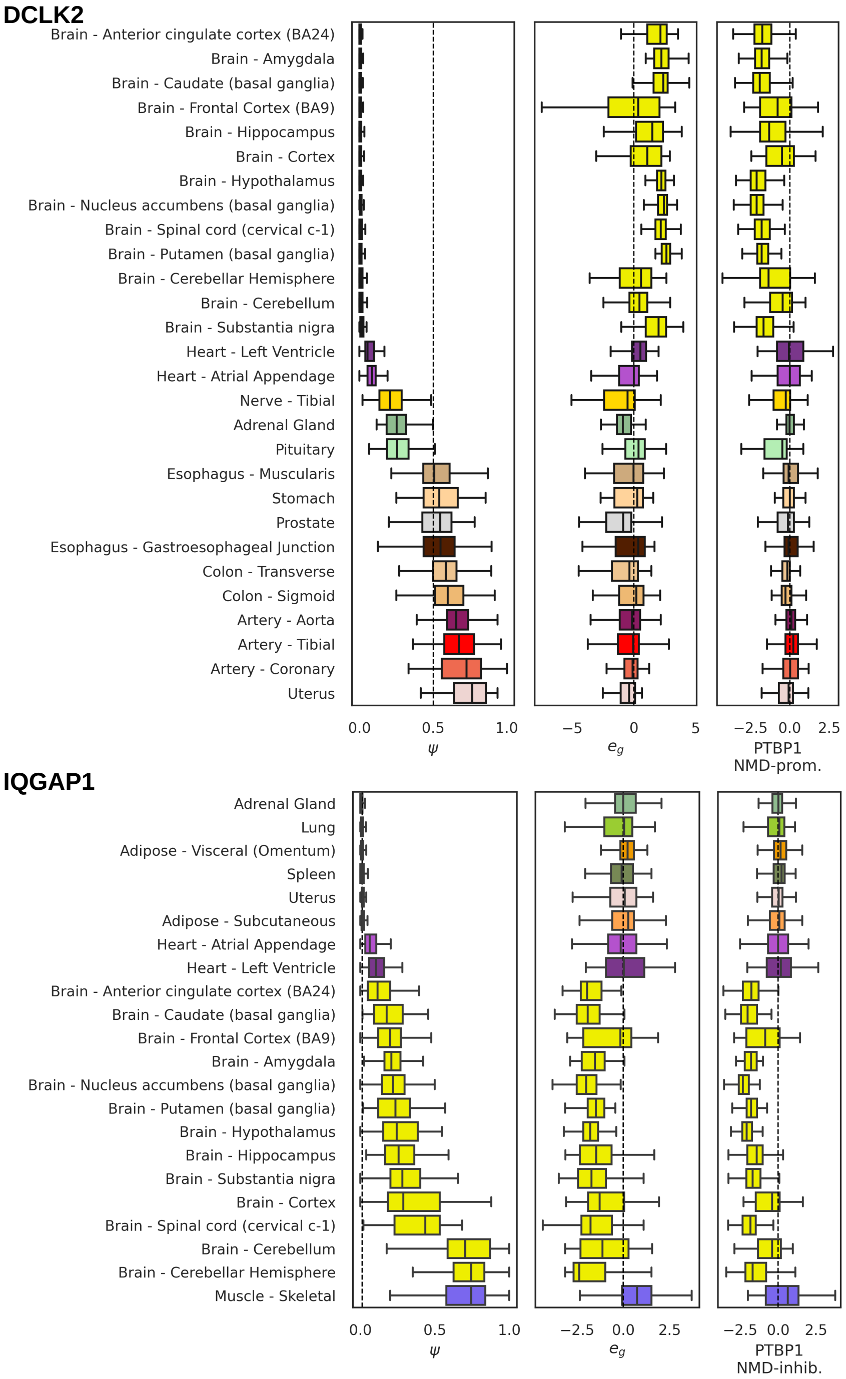

**Figure S10:** An extended diagram of NMD isoform activation and gene expression of the *DCLK2* and *IQGAP1* in tissue subtypes.

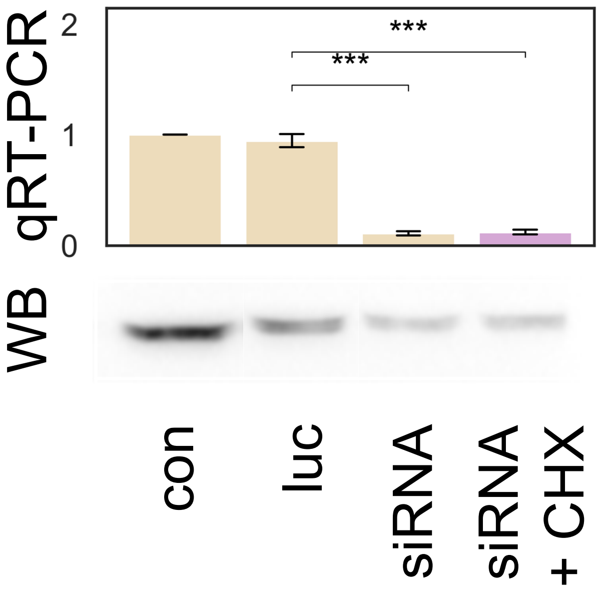

**Figure S11:** siRNA-mediated KD of *PTBP1* in A549 cell line. *PTBP1* expression was assessed by qRT-PCR (top) and Western Blot (bottom) in non-treatment control (con), upon treatment with siRNA targeting the firefly luciferase gene (luc), siRNA targeting *PTBP1*, treatment with CHX and siRNA targeting *PTBP1*.

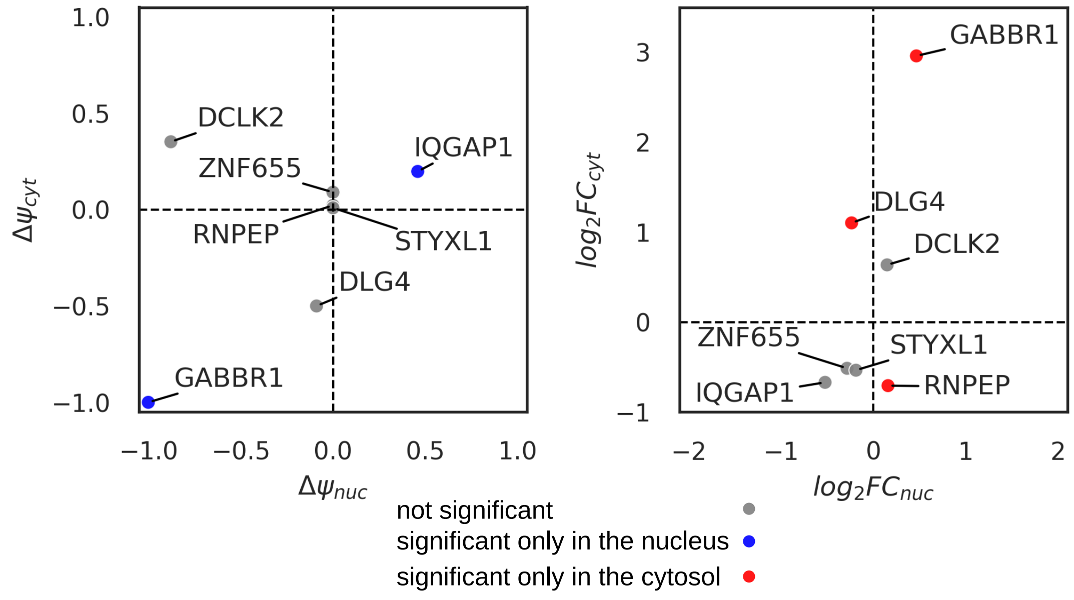

**Figure S12:** *PTBP1* KD with fractionation into cytoplasmic and nuclear RNA. The changes of $\psi$ (left) and $e_g$ (right) in response to *PTBP1* KD are shown for the predicted *PTBP1* targets in RNA-seq experiments assessing nuclear ($x$-axis) and cytoplasmic ($y$-axis) RNA fractions. Statistically significant changes (P $< 0.05$) observed exclusively in the nucleus and cytosol are marked with blue and red color, respectively. No USEs were significant in both fractions with respect to $\psi$ or $e_g$.

https://zenodo.org/record/7339827/files/ST1.xlsx

**Table S1:** The list of USEs and their regulators, for which the experimental validation was reported in human or mouse cell lines or tissue specimens.

https://zenodo.org/record/7339827/files/ST2.tsv

**Table S2:** Listed in this table are AS events that switch from a protein-coding isoform to an NMD isoform according to the GENCODE annotation (annotated USEs) and validated USEs. "PTC position" denotes the position of the PTC in the NMD isoform. The last ten columns display genomic coordinates of splice junctions or intron retention splice sites that are involved in alternative splicing.

https://zenodo.org/record/7339827/files/ST3.xlsx

**Table S3:** The list of GTEx tissues, their code and color designations.

https://zenodo.org/record/7339827/files/ST4.tsv

**Table S4:** The list of RBP perturbation experiments and their accession numbers.

https://zenodo.org/record/7339827/files/ST5.tsv

**Table S5:** The correspondence between Proteomics DB tissues and GTEx tissues (SMTS).

https://zenodo.org/record/7339827/files/ST6.xlsx

**Table S6:** siRNA sequences for the KD of *PTBP1* and primers used in RT-PCR and qRT-PCR experiments.

https://zenodo.org/record/7339827/files/ST7.tsv

**Table S7:** The classification of validated and annotated unproductive splicing events as significant and not significant. Listed are the values of $\Delta\psi$, $\Delta e_g$, $z-score$ of $\Delta e_g$, $\Delta e_l$, $z-score$ of $\Delta e_l$.

https://zenodo.org/record/7339827/files/ST8.tsv

**Table S8:** For each significant USE, shown are deviations of the $\psi$ from the pooled median, the deviation of $e_g$ and $e_l$ from the respective pooled medians. The column "Effect" indicates whether the test for tissue-specificity in a tissue gave the expected (negative) or unexpected (positive) association.

https://zenodo.org/record/7339827/files/ST9.tsv

**Table S9:** The columns "DS" and "DE" represent the changes in $\psi$ and $e_g$ under the condition of high RBP expression vs. low RBP expression, averaged over all experiments for the given RBP. NS and NT labels denote not significant and not tested cases, respectively. The last three columns list the evidence of RBP binding from CLIP data and literature citations.

`https://zenodo.org/record/7339827/files/ST10.tsv`

**Table S10:** Shown in the table are all RBP-USE pairs that were predicted from RBP perturbation assays for tissue-specific USEs. The columns “# expected” and “# unexpected” show the number of tissues with the expected and unexpected associations between $\psi$ and RBP expression. The last two columns show the evidence of RBP binding from CLIP data.

https://zenodo.org/record/7339827/files/Supplementary%20data%20file%201.pdf

**SupplementaryDataFile 1:** A gallery of 27 tissue-specifically regulated USEs with strong evidence of local RBP binding.
